# IDBac: an open-access web platform to identify bacteria and analyze relationships in culture collections using MALDI-TOF mass spectrometry

**DOI:** 10.1101/2025.10.15.682631

**Authors:** Nyssa K. Krull, Michael Strobel, Julia Saulog, Liana Zaroubi, Bruno S. Paulo, Mandisa Timba, Douglas R. Braun, Gabrielle Mingolelli, Jessia Raherisoanjato, Robert A. Shepherd, Abigail F. Scott, Carlo De Silva, Claire Fergusson, Zachary Daniel, Shailaja K. Pokharel, Sean Romanowski, Antonio Hernandez, Mónica Monge-Loría, Claire E. Dylla, Manasi M. Natu, Valentina Z. Petukhova, Neha Garg, Paul R. Jensen, Adriana Blachowicz, Chelsi D. Cassilly, Lisa Guan, D. Cole Stevens, Jaclyn M. Winter, Shaun M. K. McKinnie, Barbara I. Adaikpoh, Skylar Carlson, Erin P. McCauley, William W. Metcalf, Tim S. Bugni, Michael W. Mullowney, Eric G. Pamer, Matthew T. Henke, Hazel Barton, David O. Carter, Alessandra S. Eustáquio, Roger G. Linington, Laura M. Sanchez, Mingxun Wang, Brian T. Murphy

## Abstract

The identification and analysis of bacteria is central to the microbiological sciences. While gene sequencing methods have been the standard to achieve this, use of MALDI-TOF mass spectrometry (MS), particularly in clinical microbiology, provides high-throughput identification to the subspecies level. However, biotyping has yet to be adopted outside of clinical settings due to the lack of a centralized public database of MS protein signatures that would facilitate isolate identification via spectral comparison. Further, current platforms lack meaningful ways to compare multiple properties from large numbers of bacterial isolates. Herein we present the IDBac web platform, a crowd-sourced central knowledgebase of protein MS signatures of >1400 strains spanning 6 bacterial phyla. Accompanying the knowledgebase is analysis infrastructure to identify unknown isolates, probe relationships within culture collections using metadata integration, and visualize specialized metabolite differences within groups of closely related bacteria. To highlight this utility and encourage wide community contribution, examples of each are presented.

## Main

For over three centuries, researchers have searched for a reliable way to identify and classify bacteria. Early methods relied on morphological and metabolic features to organize isolates into distinct categories, which led to a foundational understanding of the prokaryotic domain. In the 1970’s, DNA sequencing techniques were developed to target the highly conserved 16S ribosomal RNA (rRNA) and 23S rRNA regions,^1,2^ which allowed for bacterial identification based on a single gene. The development of polymerase chain reaction (PCR) and incorporation of automation in sequencing methods allowed for the first full bacterial genome sequences in 1995.^3–6^ Currently, affordable sequencing costs and increased access to bioinformatic platforms have resulted in greater than 400,000 genomes available in public databases and catalyzed a new era of understanding in bacterial diversity, phylogenetic classification, and overall bacterial function.^7^ However, it is not yet practical for whole genome sequencing (WGS) to be employed as a means to rapidly identify and organize large numbers of unknown bacterial isolates.^8^

In 1996, matrix assisted laser desorption/ionization time of flight mass spectrometry (MALDI-TOF MS) was demonstrated to be an orthogonal approach to identify bacteria.^9,10^ Briefly, bacterial cell mass or extract was ionized and the resulting spectrum of intact high copy number protein signals acted as a distinct fingerprint and proxy for its phylogenetic identity. The most widespread application of this technique has been in clinical laboratories.^11–14^ Since 2009, a growing number of companies such as Bruker and bioMérieux have compiled and commercialized protein MS reference databases of clinical pathogens and designed proprietary software for strain identification, which in some cases provides subspecies-level strain resolution.^15–21^ While online search services such as MicrobeNet^22^ exist, they are also tailored toward clinical libraries and require vendor-specific file types, which limits open access and cross-vendor usability. Further, MicrobeNet database search results are emailed directly to users and broader analysis including dimensionality reduction, clustering, and metabolite production is unavailable. MicrobeMS is a longstanding, effective resource that adopts the open.mzXML format and expands beyond identification to additional statistical analyses (e.g., hierarchical clustering and biomarker peak analysis), mitigating several of these challenges.^23,24^ Nonetheless, the lack of a central spectral repository that includes environmental microorganisms remains a critical limitation that prevents open sharing of spectral data. A comparison of some available databases and analysis platforms is included in **Supplementary Table S1**.

To address barriers imposed by the proprietary nature, cost, and emphasis on pathogenic microorganisms that render existing resources impractical for broader usage, we introduce the IDBac web platform. The IDBac web platform fulfills the need for a central community-driven MALDI-TOF MS knowledgebase and accompanying analysis platform to identify and compare bacterial isolates, especially those collected from environmental or host-derived niches. Numerous review articles and perspectives spanning greater than a decade have noted this as a major limitation of the MALDI MS approach and have called for the creation of a public central repository of protein MS signatures.^11,25–32^ Here, we present three major analysis features of the platform.

### Creation of an open access, community-sourced database of bacterial protein MS profiles

Several research groups have released MALDI-TOF protein MS spectra of characterized bacterial isolates, though these were not designed to be a lasting community resource. Many of these efforts employed proprietary data formats and were limited in scope (i.e., fewer than 100 strains from a single bacterial genus or species). ^26,33,34^ Notable efforts include SPECLUST,^35^ PVBase,^36^ VibrioBase,^37^ collections from NASA Jet Propulsion Laboratory ^16,38^ and the Drinking Water Library,^33^ and collections of highly pathogenic bacteria.^23^ However, these data are not associated with a unified public platform that allows spectra to be compiled, compared, and analyzed. To address these limitations, we introduce the IDBac web platform, an open-access knowledgebase (IDBac-KB) combined with web accessible computational analysis pipelines (IDBac-Analysis). To date, the IDBac-KB provides access to MS data from 1,418 bacteria spanning 151 genera and 6 phyla and allows users to upload protein MS spectra for community use. IDBac-Analysis enables users to query protein MS profiles against the IDBac-KB to putatively identify ‘unknowns’; view strain/isolate protein MS similarity groupings in bulk via dendrogram analysis; integrate experimental metadata within a dendrogram to explore trends in strain/isolate collections; and when data are also collected in reflectron mode, visualize specialized metabolite differences within groups of closely related bacteria (**Fig. 1A-B**). Through several examples described in this manuscript, we show how IDBac-Analysis can be used as a hypothesis generation tool *in the front end* of a researcher’s analysis pipeline.

**Fig. 1.**
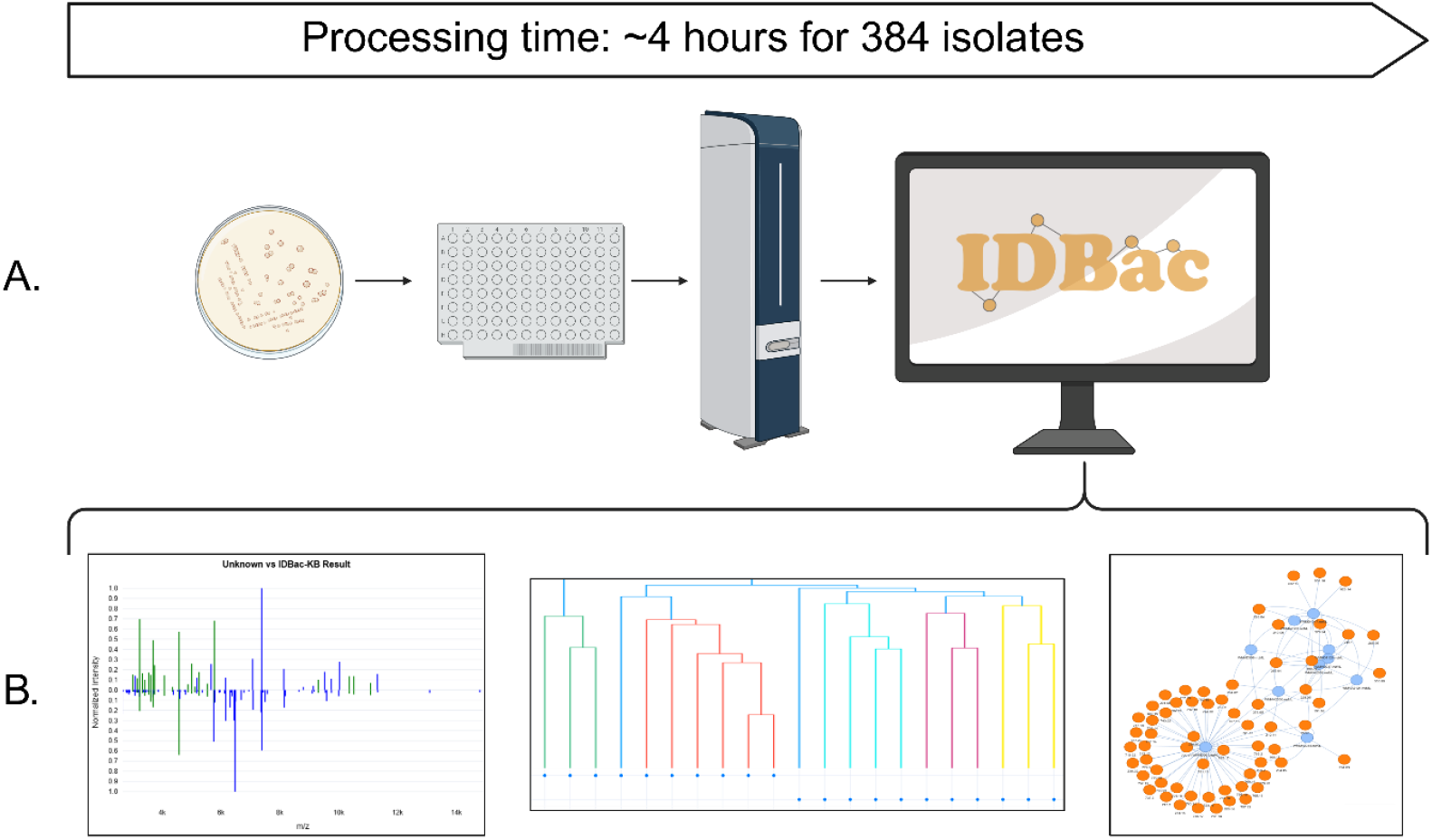
The IDBac web platform offers a cost effective and high throughput approach to analyze MALDI-TOF MS data. **A**. Cell material from single bacterial colonies are transferred to a MALDI-TOF target plate, data are acquired, and spectra files are uploaded to the IDBac/GNPS2 interface for data processing. **B**. The IDBac web platform offers putative identification using the IDBac-KB (Feature 1), multiple protein MS data visualization tools (Feature 2), and metabolite association networks for specialized metabolite assessment (Feature 3). Of note, this time estimate was made using an axial MALDI-TOF that can acquire both linear and reflectron mode data with a maximum laser frequency of 2 KHz. The time estimate represented in this figure would change with access to different instrumentation. Figure created in BioRender. Krull, N. (2026) https://BioRender.com/697lx8t

## Results

### IDBac-KB, a public knowledgebase of bacterial protein MS signatures

To bootstrap IDBac-KB, we collected MALDI-TOF MS spectra of strains from in-house collections and worked with collaborators to both import their data and analyze their collections. In-house data collection was performed on Bruker Microflex, Autoflex, and RapifleX with a 1,800-21,000 Da mass range in biological replicates varying between 3 to 8, with 1,500-2,000 shots acquired per well (See Methods: MALDI-TOF MS protein data collection). When importing into the IDBac-KB, replicate MALDI MS spectra were merged into a single consensus spectrum per isolate and manually inspected for quality control. Metadata such as Genbank accessions, 16S rRNA sequences, source, isolation, and growth conditions associated with each strain were compiled and deposited into the knowledgebase. External identifiers such as Genbank accessions are integrated with their respective resources to enrich the knowledgebase with detailed and up-to-date taxonomic information (**Supplementary File 1**).

IDBac represents a substantial contribution to open-access MALDI-TOF MS protein spectral repositories. At the time of publication it consists of 1,418 entries that comprise 6 bacterial phyla, 72 families, 151 genera and 472 species (**Fig. 2**) and to our knowledge is the largest open-standard format spectral database to date. These numbers exclude public datasets that were incorporated into the KB. The integration of version 4.2 of the Robert Koch Institute Database of Highly Pathogenic Microorganisms^23,39^ substantially augmented our efforts and increased coverage to 3,399 entries, 8 phyla, and 166 genera. We note, however, that analysis/results presented in this manuscript do not include strains from the Koch database. Of the 1,418 strains in the KB, 283 have been characterized by whole genome sequencing (WGS) and 1,135 by 16S rRNA gene sequencing analysis. Strains must have associated 16S rRNA data at the minimum to be accepted into the KB, however unknown isolates are accepted for the IDBac-Analysis pipelines. The database received major contributions from government institutions such as the NASA Jet Propulsion Laboratory and Marshall Space Flight Center (111 strains, 30 genera); government-academia partnerships that contributed a portion of the United States Department of Agriculture (USDA) NRRL culture collection (420 strains, 9 genera); academic institutions such as the Duchossois Family Institute microbiome culture collection (156 strains, 15 genera); and 16 academic laboratories that contributed their culture collections (620 strains, 127 genera). Collectively these allow for a massive expansion of available reference spectra and provide a community resource with rich taxonomic and geographic diversity.

**Fig. 2.**
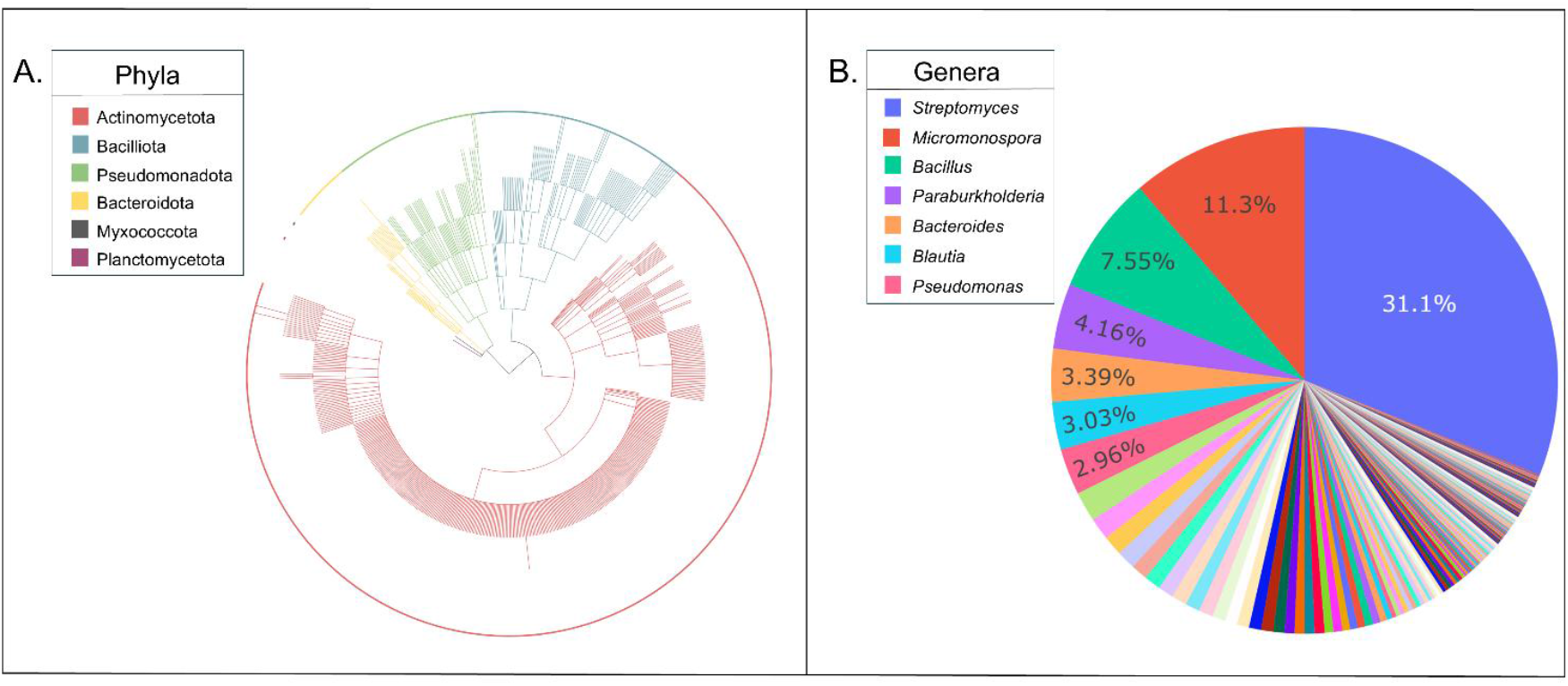
The IDBac protein MS database hosts diverse bacterial taxa. **A**. Taxonomic tree representing current strain entries in the database (1,418 entries comprise 6 bacterial phyla, 151 genera and 472 species) created using NCBI hierarchical classifications. **B**. Database composition color-coded by genus, legend lists genera with >30 strain representatives.

### Feature 1: Querying MS protein spectra from bacterial isolates against IDBac-KB

IDBac-Analysis provides putative bacterial identities to user-submitted protein MS spectra that are measured from either a small amount of cell material or extracted cell mass. Query spectra are processed according to **Methods: IDBac-Analysis Workflow** and searched against protein MS spectra of genetically characterized strains within the IDBac-KB using an intensity-agnostic cosine distance score. KB “hits” are then ranked by cosine distance (on a 0-1 scale, 0 signifies identical), which provides a measure of spectral similarity that reflects taxonomic relatedness between strains.^40^ Results from querying the MALDI MS spectrum of a single environmental isolate collected from the surface of a marine snail are depicted in **Fig. 3A** and **Supplementary Fig. S1**. Environmental isolate Z001 most closely matched to a *Solwaraspora* sp. from a collection at the University of Wisconsin Madison (top hit in the IDBac-KB with a cosine distance of 0.67). WGS sequencing identified Z001 as a *Solwaraspora* sp., confirming the genus assignment provided by IDBac with an Average Nucleotide Identity (ANI) of 79.9 using fastANI.^41^ We note that when using cosine distance to evaluate protein MALDI MS spectral similarity, there is no universal threshold to define genus/species boundaries between taxa. Since this value must be determined on an individual basis, IDBac reports the top hits for each query along with a relatively relaxed user-adjustable threshold for cosine distance (0.7). A better understanding and defining of these boundaries is an ongoing challenge in the field of bacterial mass spectrometry.^42,43^

**Fig. 3.**
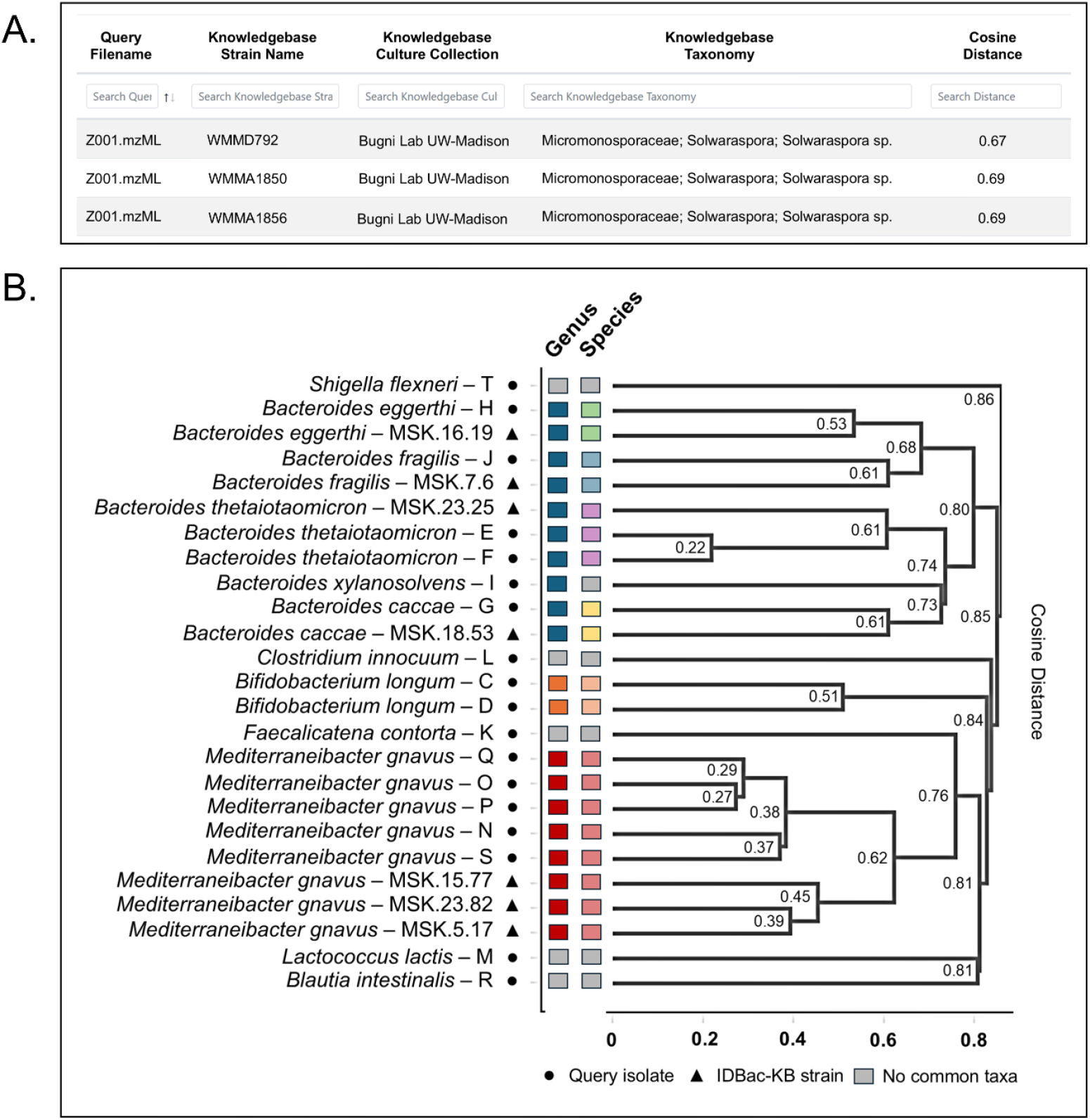
The IDBac-KB allows for rapid identification of unknown isolates. **A**. A knowledgebase search summary lists cosine distances between the query isolate (Z001) and the top three KB hits alongside strain, culture collection, and taxonomy. **B**. A protein MS dendrogram displays genus and species-level discrimination between isolates from two distinct bacterial microbiome collections and highlights the ability of IDBac to obtain identifications for multiple isolates simultaneously. Colors highlight common genera and species within the dendrogram. Note: there is currently no universal threshold to define genus/species boundaries between taxa; this is a highly underdeveloped research field.

IDBac-Analysis supports high-throughput taxonomic annotation with batch MALDI MS acquisitions. As a demonstration, we performed a blind analysis of 18 in-house isolates from the human gut microbiome. Consensus spectra (representing all replicates of each isolate) were simultaneously queried against the IDBac-KB using a cosine distance threshold of 0.65 (see **Methods:** MALDI-TOF MS target plate preparation and IDBac Analysis Workflow). This search required less than five minutes and resulted in the putative identification of ten isolates with high similarity to IDBac-KB strains contributed by the Duchossois Family Institute at the University of Chicago, who maintains a large collection of human gut microbiome strains. 16S rRNA gene sequencing analysis confirmed the genus and species assignments provided by IDBac (**Supplementary Table S2**).

Additionally, IDBac-Analysis can visualize taxonomic diversity with hierarchical clustering, where a protein MS-based dendrogram groups isolates based on computed distances of protein MS fingerprint similarity (**Fig. 3B**). As expected under average-linkage clustering, the ten microbiome query isolates clustered with their corresponding IDBac-KB hits. These human gut-associated isolates were annotated with 16S rRNA gene sequencing and revealed broad agreement between phylogenetic and proteomic relationships inferred from MALDI data. Specifically, at a cut height of 0.74 in the dendrogram, all clusters contained exactly one genus. Notably, intra-species matches dominate the lowest branch distances (e.g., *Bacteroides thetaiotaomicron* queries match at a height of 0.22). However, it is commonly observed that different species within a genus can be split into multiple groupings across the dendrogram. This is a common phenomenon in MALDI MS dendrogram analysis and highlights a contrast with WGS-based phylogenetic trees: while proximity in the protein MS dendrogram serves as information that two strains are likely related, all species within a genus are not necessarily grouped below an assigned cut height. This finding is consistent with the knowledgebase search results that show while the highest precision is achieved at lower thresholds, additional matches within a genus may occur at higher distances (**Supplementary Figure S2, Table S3**).

### Feature 2: Integration of isolate metadata into IDBac dendrograms

The incorporation of metadata that details a variety of experimental and environmental conditions into analyses is common in microbiological sciences.^44–48^ Further, the integration of metadata to enhance large metabolomics analyses has been performed, though cross-user comparisons remain a challenge in the field for a variety of reasons.^49^ However, this practice is often done post-hoc and its manual integration remains a major roadblock for its successful application at the repository scale. IDBac-Analysis has significantly expanded past efforts to organize MS data from bacterial isolates into dendrograms that are enriched with metadata context.^50–53^ Users can now observe trends in metadata from large collections of bacteria when visualized as a dendrogram, protein intensity heatmap (**Supplementary Fig. S3**), or dimensionality reduction plots. To demonstrate these features, we first analyzed 79 isolates collected from a lake within a cave system in South Dakota. Hierarchical clustering revealed homophilic patterns in which samples from within source (lake/sediment) clustered closely (**Methods**). Additional statistical analysis revealed significant differences in dispersion (PERMANOVA: pseudo-F=9.633738, p=0.001, n=79,999 permutations; PERMDISP: F=10.55, p=0.001) and Principal Coordinates Analysis (PCoA) indicated a shift in group centroids between sample sources. Additionally, 16S annotations were integrated highlighting differential genus annotations between sample sources (lake/sediment) in addition to divergence in proteomic profiles within genera (**Supplementary Note 1A**). Specifically, these data provide evidence that subclades – driven by MALDI MS protein homology – were only observed in specific regions within the cave, i.e. sediment versus lake water (**Fig. 4A**). This insight is possible due to analysis of the metadata-integrated dendrogram and has potential to offer valuable insight into relationships between environmental samples and their corresponding bacterial consortia.

**Fig. 4.**
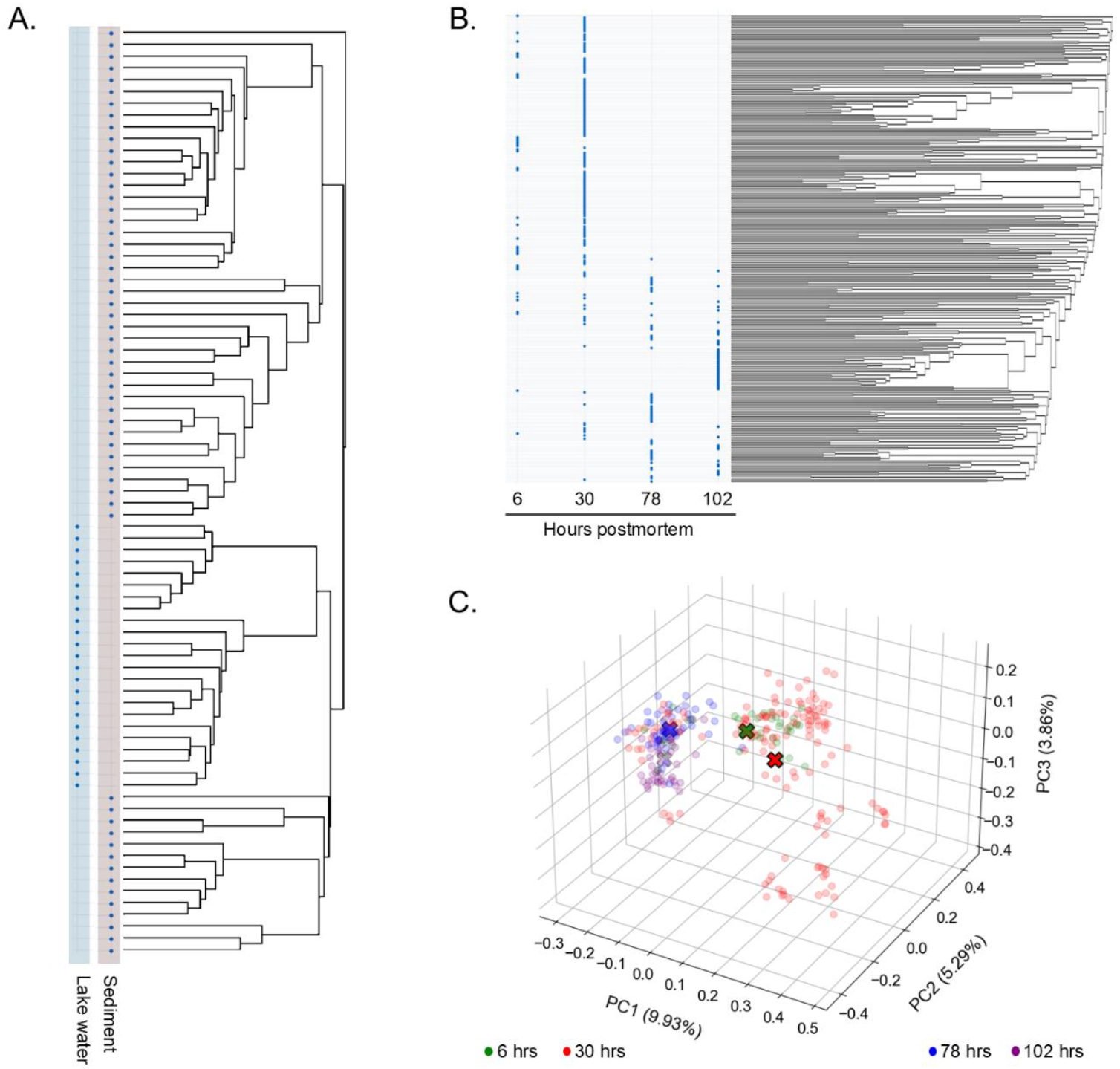
Metadata-integrated protein MS dendrogram visualizes trends within culture collections. **A**. Metadata enrichment shows cultured bacterial populations are non-overlapping between two sample types collected from Wind Cave, SD. **B**. Metadata analysis reveals taxa-specific trends as a function of hours postmortem during the decomposition of a pig carcass. **C**. Principal Coordinates Analysis (PCoA) shows multivariate MALDI spectral profiles of each pig carcass isolate, colored by their source timepoint. ‘X’ symbols represent timepoint centroids. Statistically significant differences in distributions between timepoints is observed (PERMANOVA=0.001, PERMDISP=0.001). The PCoA visually confirms a centroid shift and differences in dispersion with the largest shift between successive timepoints was observed along principal component 1 (PC1) between the 6-hour centroid (PC1: 0.0295) and the 30-hour centroid (PC1: 0.1294). Overall, these findings show the trajectory of spectral profiles over time as bacterial communities change during carcass decomposition.

A second example highlights work in the field of forensic taphonomy, where an association between postmortem intervals and pig cadaver microbial populations was investigated. In this study, skin swabs of a pig carcass were collected at consecutive time points over the course of 4 days, resulting in 274 axenic bacterial cultures (**Methods**).^54^ Each isolate was processed as described in **Methods** and MALDI MS data in the protein mass range were collected and visualized as a dendrogram using IDBac-Analysis. Sample timepoint metadata was included and the visualization suggested distinct populations of taxa were present at different timepoints, providing preliminary evidence for bacterial variation across different states of decomposition (**Fig. 4B**). A significant difference in dispersion was observed (PERMANOVA: pseudo-F = 8.47, p = 0.001, n = 273, 999 permutations; PERMDISP: F = 220.73, p = 0.001) and PCoA was performed, revealing a shift in proteomic centroids between collection time points. (**Fig. 4C, Supplementary Note 1A-1B**). These results are consistent with previous work that documented an association between bacterial diversity and insect activity across postmortem decomposition stages.^55^ Insect occupation occurs within the initial 6 hour period (with minimal decomposition), followed by a network of decomposing microorganisms by 30 hours postmortem. A shift in community is again observed in later stages of decomposition, which aligns with reported depletion of cadaver-associated nutrients and moisture content. Minimal overlap in the IDBac dendrogram across sampling periods highlights these published trends and offers an orthogonal approach to estimating postmortem interval. To achieve similar results using traditional bacterial identification methods such as 16S rRNA gene sequencing, sample preparation of 274 isolates and manual curation of metadata would require a minimum of one week of benchwork and cost thousands of dollars, further emphasizing MALDI-IDBac analysis as a cost-effective and high throughput alternative.

### Feature 3: Visualization and analysis of Metabolite Association Networks (MANs) in IDBac-Analysis

IDBac-Analysis also extends to the analysis of specialized metabolites (200-2,000 Da), a dataset that can be acquired consecutively from MALDI instruments. This is typically acquired in reflectron mode although acquisition can also be carried out in linear mode depending on user access to specific types of instrumentation. Incorporation of this analysis can enhance bacterial differentiation and increase accessibility of specialized metabolite production to non-experts. It is worth noting that only protein analysis is conducted while biotyping in clinical microbiology labs, which differentiates our work from current clinical practices. This concept has proven to be an effective means to discriminate between bacterial isolates at the subspecies level even when isolates share nearly identical phenotypic traits or 16S rRNA gene sequences.^56^ The analysis workflow visualizes relationships between bacterial isolates by displaying both conserved and unique masses (representing the specialized metabolites) between different strains. Subtraction of a matrix blank is employed to remove *m/z* signals attributable to the matrix. Available as an interactive online dashboard, MANs in IDBac-Analysis were designed for and are accessible to a wide audience without extensive background in computational mass spectrometry. One key analysis approach is to assess patterns of specialized metabolite production as a function of phylogenetic relatedness, which can provide a means to discriminate strains at the subspecies level. We note, however, that while MS1-based data provide a signal for differential metabolite expression, they do not discriminate between isobaric compounds and are inherently limited by an instrument’s mass resolving power (see **Supplementary Note 2**).

### Bacterial species discrimination and new compound discovery using comparative MANs

As part of an effort to mine a collection of Burkholderiales strains for novel specialized metabolites, a small amount of cell material from 316 strains was analyzed for both protein and specialized metabolite spectra. In this collection, we identified two *Paraburkholderia* strains that were indiscernible through 16S rRNA gene sequencing analysis (1-D8 and 3-E9, 100% pairwise identity, 1,473 bp). BLAST analyses of the 16S rRNA sequences classified both 1-D8 and 3-E9 as *Paraburkholderia megapolitana*. However, using IDBac-Analysis, we observed differential metabolite production profiles via MAN analysis (**Fig 5A**). Subsequent WGS analysis revealed significant differences between the genomes and these bacteria were reclassified as *Paraburkholderia megapolitana* (3-E9) and *Paraburkholderia acidicola* (1-D8).^57^ Importantly, early in the pipeline IDBac MAN analysis indicated that two seemingly identical strains grown under the same cultivation conditions offered divergent specialized metabolite production, which in this case was an indicator for species-level differentiation. This highlights the power of metabolomics to sometimes act as a cost-effective indicator of phylogenetic divergence and alleviates some of the cost burden of WGS, particularly when large collections of bacteria are involved.

**Fig. 5.**
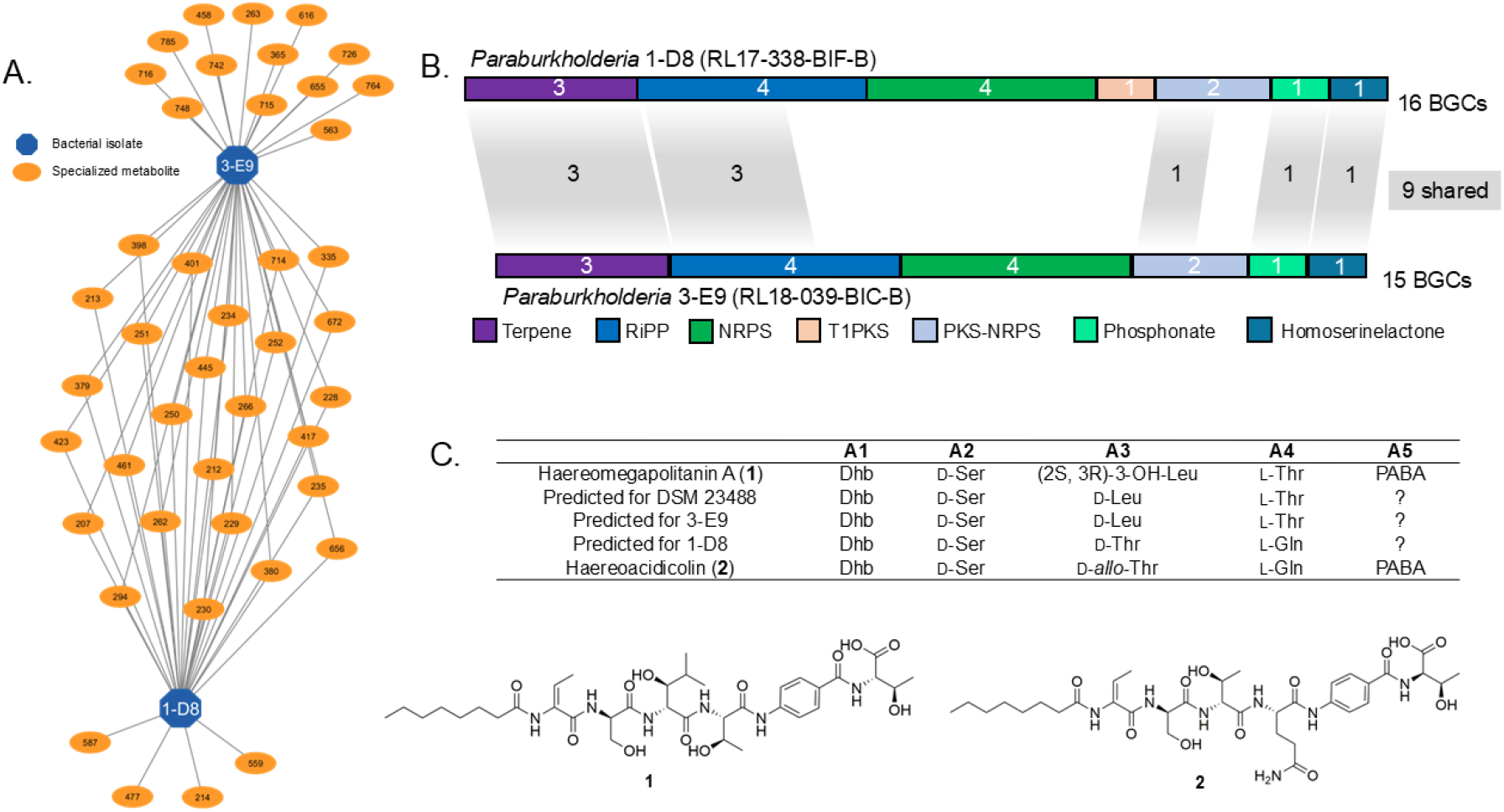
IDBac can discern seemingly identical strains. **A**. IDBac MAN of strains 3-E9 and 1-D8. **B**. Genome data. Comparison of shared and unique BGCs between strains 1-D8 and 3-E9 (shaded lines). **C**. Structure of haereomegapolitanin A (**1**) detected in strain 3-E9 and previously reported from DSM 23488. Strain 1-D8 contains a related gene cluster but with distinct amino acid predictions based on the A and C domains in the NRPS (as shown in the table) and produces haereoacidicolin (**2**).

Comparative biosynthetic gene cluster (BGC) analysis on antiSMASH highlights these differences. Strain 3-E9 and 1-D8 possessed 15 and 16 BGCs respectively, 9 of which were shared (**Fig. 5B**). Of particular interest were two related nonribosomal peptide synthetase (NRPS) BGCs, one of which (strain 3-E9) aligned closely with the published BGC for haereomegapolitanin A.^57^ Examination of UPLC-MS/MS data for the 3-E9 extract identified a metabolite with high agreement to the published MS/MS spectrum for haereomegapolitanin A (**Supplementary Fig. S4**). In contrast to strain 3-E9, *Paraburkholderia* 1-D8 possesses a related gene cluster with variations in the domains that dictate amino acids A3 and A4. Bioinformatic analysis of adenylation and condensation domains suggested the replacement of hydroxy-leucine with D-threonine (A3) and L-threonine with L-glutamine (A4). Large-scale fermentation revealed the presence of a new metabolite with the predicted protonated molecule (calc. 764.3825, obsd. 764.3858, 4.32 ppm). Isolation and structure elucidation using a combination of high-field NMR and high-resolution mass spectrometry established the planar structure of this new metabolite (**Fig. 5C, Supplementary Fig. S5 & S6-15, Supplementary Note 3, Supplementary Table S4**). Marfey’s analysis supported by bioinformatics analysis of adenylation and condensation domains confirmed the predicted configurations at all chiral centers in the molecule (**Supplementary Note 3, Supplementary Fig. S16-19**), and the compound was named haereoacidicolin (**2**).

This discovery highlights the shortcomings of 16S rRNA sequencing to distinguish bacterial strains. It also stresses that although WGS remains an effective way to identify and detail differences between phylogenetically similar bacteria, it is not always a cost-effective means to screen for differences in a large collection of isolates. Here, IDBac-Analysis of MANs was used to prioritize two seemingly identical strains from a collection of 316 for a more in-depth metabolic investigation, where WGS then detailed the biosynthetic differences and the new metabolite was purified and characterized. It is important to note that the MANs provide a direct measure of expression for the metabolites, whereas genome level predictions cannot anticipate the active transcription of a given gene.

## Discussion

The IDBac Platform facilitates the use of bioinformatics tools and offers users the freedom to run analyses anywhere, on most browser-enabled devices. The original desktop software version of IDBac (that did not host a KB or several other functions described herein) saw strong community engagement with >1,500 downloads worldwide. The current IDBac web platform is receiving a similar response with >2,479 unique site visits spanning 60 countries, pre-publication. This community-centered resource offers bacterial identification and a suite of customized analyses applicable across the broad microbiological sciences. The creation of the IDBac-KB addresses a long-standing appeal for a taxonomically diverse, open-access protein MS repository that bridges the accessibility gap between for-profit resources and the broad research community. Just as the Basic Local Alignment Search Tool (BLAST)^58^ expanded the capacity of researchers to identify unknown bacteria in the lab by providing access to crowdsourced nucleotide sequences, IDBac is anticipated to function similarly via community contributions of crowdsourced MS protein spectra.

IDBac-Analysis has the capacity to provide proteomic and metabolomic analyses from a small amount of bacterial cell mass in a comparatively short measure of time. The resulting data provide a unique opportunity to observe relationships between large collections of unknown isolates and to empower researchers to form hypotheses, particularly near or at the beginning of a research project. Recent examples include the dereplication of phylogenetic and metabolomic redundancy from a large culture collection,^52^ exploration of the relationship of bacterial phylotype with specialized metabolite production in aquatic hosts,^42^ and selective mining of taxa from the environment.^43^

However, there are several notable limitations associated with any crowd-sourced repository. First, the health of the database is reliant on users conducting quality control (QC) of their spectra prior to deposition. Inconsistencies in data quality could diminish spectral matching over time, resulting in inaccurate putative identities. To address this, IDBac offers a ‘dry run’ deposition with integrated spectra visualization that allows users to manually inspect spectra before contributing to the public-facing database. Additionally, we have provided detailed guidance on best practices for MALDI-TOF MS sample preparation and QC (see **Data Availability**). Second, the efficacy of this method to identify bacteria is dependent on having a large and taxonomically comprehensive reference dataset. For example, some genera such as *Bacillus, Micromonospora*, and *Streptomyces* harbor a diverse array of protein MS fingerprints that cause a wide distribution of its species across a dendrogram, as these groupings are driven by more than just ionized proteins of ribosomal origin. A given *Streptomyces* isolate may show low homology to the IDBac-KB despite there being many species represented. Therefore, contributions from the global scientific community are paramount for the development of a robust protein MS repository that reflects the diverse and expansive bacterial domain.

Third, inclusion of thorough metadata is a crucial and often overlooked aspect of user-contributed databases. Lack of transparent and detailed protocols undermines confidence in data quality and hinders reproducibility, particularly when data are sourced from biological materials. The IDBac database requires select metadata categories for successful depositions, however users are encouraged to provide additional information associated with collections, not only for the sake of data availability but for the potential of these data to advance the field through unanticipated future needs. To aid in these efforts users are encouraged to provide information for all categories listed within the provided metadata template (**Supplementary File 2**). To further highlight the utility of these metadata, a retrospective analysis was conducted showcasing the efficacy and reproducibility of bacterial identification using the IDBac platform across laboratories (**Supplementary Notes 4** and **5, Supplementary Fig. S20-22**).

Fourth, currently the mechanism of grouping isolates within IDBac is limited to three distance functions (Cosine, Presence, Euclidean), which is a subset of existing literature. There is no distance function that is universally most effective at grouping bacterial isolates; each function has advantages and limitations. We caution, however, that spectral distance is both a function of the distance function itself and how the raw spectra are processed into peak lists. Therefore, to the best of one’s ability each experimental dataset should be scrutinized accordingly. Additional distance metrics will be tested as the database continues to expand.

Finally, there is often a perception that MALDI MS is a specialized instrument that lags other platforms in mass resolving power and is only meant for proteins due to interference or ion suppression from the matrix peaks themselves. The protein spectral analyses rely on the groundwork established by the FDA approved biotyping protocol and we have shown that the resolution of peaks for both protein and specialized metabolite spectra greatly aid in dereplication of microbial isolates. Despite the limitations noted in **Supplementary Note 2**, we have shown that unit-resolution peaks from specialized metabolite spectra greatly aid in dereplication of microbial isolates. Future users are advised to inquire with their core facilities or clinical microbiology labs for access to MALDI TOF instrumentation. Additionally, while MS1 data are currently accepted as input for MANs, IDBac-Analysis is not designed to provide putative compound identifications. The IDBac pipeline is being expanded to include MS/MS specialized metabolite data such as that from QqTOF or Orbitrap-based instrumentation in the near future. IDBac is flexible and enables analyses from a variety of instruments and mass analyzers, which expands the accessibility of this approach.

## Methods

### Strain acquisition, incubation, and MALDI-TOF MS sample preparation

To create an initial database population, strains whose identity was determined through WGS or 16S rRNA gene sequencing were obtained through collaboration with academic laboratories and governmental institutions (**Supplementary Table S5**). IDBac-KB entries were also sourced from in-house strain collections. Prior to MALDI-TOF MS data collection, the majority of strains used in this study were incubated on A1 agar media at ambient temperature for 5-7 days, with exceptions for fastidious organisms or those requiring extended incubation time. Specific cultivation conditions and sequencing data for each database entry can be found within the IDBac-KB (**https://idbac.org/database**).

Isolates used in the decomposition study were collected from skin swabs of a domestic pig carcass (*Sus scrofa domesticus*) taken at various time points and stored at −20 °C until the time of plating (note: no animals were killed for the purpose of this study). Four swabs (6 hrs, 30 hrs, 78 hrs and 102 hrs) were directly applied to three types of solid substrate media containing cycloheximide (28 μM) to inhibit fungal growth, resulting in 12 bacterial diversity plates. Each discernable colony was then transferred to A1 agar media and incubated at ambient temperature for 5-7 days, yielding 273 axenic cultures, which were processed for MALDI-TOF MS data collection as described below. All carcasses were acquired commercially (Island Farms, Waianae, USA) and handled in accordance with State of Hawaii Statute §159–21. For environmental sampling details and media ingredients, see **Supplementary Tables S6** and **S7**.

Cave system isolates were derived from two sediment samples and two water samples collected from a lake in Wind Cave (Wind Cave National Park, South Dakota). The water and sediment samples were collected using a 10 mL sterile syringe: the water samples were collected ∼10 cm below the surface of the lake, while the syringe was used as a vacuum cleaner to suck sediment from the bottom of the lake. The 10 mL of sample was then injected into a glass vial containing three types of broth media supplemented with cycloheximide (28 μM; to prevent the growth of fungi). The lakes contain a large percentage of ultrasmall (<0.2 μL) cells.^59^ To enrich for these species, a sterile 0.2 μm Millipore PTFE membrane filter was placed in-line with the needle when the sample was injected into the vial. After approximately eight weeks of incubation at ambient temperature, 100 μL of each sample-vial was aliquoted onto solid substrates of their respective media and incubated at ambient temperature for two to three weeks until colony formation was observed. Each colony was isolated on A1 agar and processed for protein MS data collection as described below. For media ingredients see **Supplementary Table S8**. Samples used in these studies were collected under permit from locations described above. All other strains used for the purpose of this study were sourced from preexisting strain collections.

### MALDI-TOF MS target plate preparation

Following incubation, a small amount of cell material was transferred to wells of a ground-steel MALDI-TOF MS target plate using sterile toothpicks, in biological replicates of between 3-8. After application of colony cell mass, 1 μL of 70% formic acid (7:3 Optima, Fisher Chemical: Optima LC-MS Grade Water Fisher Chemical) was aliquoted per well. Following evaporation, an overlay of 1 μL of α-cyano-4-hydroxycinnamic acid matrix (α-cyano-4-hydroxycinnamic acid powder [98% pure, Sigma-Aldrich, part-C2020], 50% acetonitrile, 47.5% water, and 2.5% trifluoroacetic acid) was applied.

### MALDI-TOF MS data collection

MALDI-TOF MS data were collected on the following instruments: Autoflex Speed LRF mass spectrometer (Bruker Daltonics), rapifleX MALDI Tissuetyper mass spectrometer (Bruker Daltonics), Microflex LT MALDI-TOF mass spectrometer (Bruker Daltonics), and MALDI-8020 Benchtop mass spectrometer (Shimadzu). Instrument specifications for initial IDBac-KB populations can be found in **Supplementary Table S9**. Protein spectra were collected in positive linear mode and externally calibrated with Bruker Daltonics Bacterial Test Standard (BTS) or Protein Calibration Standard 1. Specialized metabolite spectra were collected in positive reflectron mode and externally calibrated with Bruker Peptide Calibration standard.

### Quality control of IDBac database submissions

Manual quality control analysis was conducted on each entry deposited in the protein MS database. Consensus spectra were assessed for intensity, signal to noise, number of replicates and peak shape. Strains possessing fewer than 3 replicates, or that failed the previously mentioned parameters were omitted from the database (See **Supplementary Fig. S23** for examples).

### 16S rRNA gene sequencing preparation and analysis

DNA was extracted from single bacterial colonies using a modified heat-lysis protocol.^60^ Pure cultures were incubated in 100 µL of 1x IDTE pH 8.0 for 5 minutes at 90 °C, followed by centrifugation at 10,000 RPM for 3 minutes. 3 μL of the supernatant was used at template DNA and transferred to a 0.2 mL Eppendorf tube containing: 8.5 μL molecular grade water, 12.5 μL KAPA2G Robust HotStart ReadyMix, and 0.5 μL of 16S rRNA universal primers 27F (5′-AGA GTT TGA TCC TGG CTC AG--3′) and 1492R (5′-GGT TAC CTT GTT ACG ACT T-3′). PCR conditions included a primary denaturation at 95 °C for 5 minutes then 35 cycles of denaturation at 95 °C for 30 seconds, annealing at 60 °C for 30 seconds, extension at 72 °C for 1 minute with a final elongation at 72 °C for 5 minutes. PCR product purification and Sanger sequencing were performed by Eurofins Genomics (eurofinsgenomics.com/en/products/dna-sequencing). Pairwise alignment, generation of consensus sequences and Basic Local Alignment Search Tool (BLAST) analysis were conducted using Geneious Prime software Version 2023.2.1. Taxonomies were assigned based on >98% identity and pairwise alignment with strains in the NCBI Genbank database.

### GNPS2/IDBac workflow methods

The full IDBac software suite consists of a database deposition and analysis set of workflows in addition to an interactive analysis page and database that we describe in the following section. The Nextflow ^61^ workflows are hosted on the GNPS2 web platform and allow users to analyze and deposit data through a graphical user interface.

### IDBac-Analysis Workflow

The IDBac-Analysis workflow enables users to query and organize MALDI-TOF spectra of bacteria isolates against the IDBac-KB. Using the MALDIquant R package^62^ protein spectra are square root transformed, smoothed with the Savitzky Golay^63^ filter (half window size=20), and baseline corrected with 100 iterations of the TopHat method. Noise is detected and removed via the mean absolute deviation method using a half-window size of 20 Daltons and a signal to noise ratio of 4.^52^ Each file is checked for formatting errors, flat-line spectra, and failed uploads. Errors and warnings are displayed to the user in the graphical user interface.

To compute database distances, protein spectra are discretized into 10 Dalton bins, intensities are normalized with the base peak set to 1.0 and averaged across replicates. Binned peaks present in fewer than 50% of replicates are removed to comprise a consensus spectrum for a given isolate. Mass ranges are truncated to a user-specified window. Using a specified distance function (with options being cosine and Euclidean distance), pairwise distance calculation is performed between all pairs of user-uploaded spectra and database MALDI MS spectra. The “Presence” option computes the cosine distance while ignoring peak intensities that may help mitigate the effects of intensity variation between runs.

In addition, the workflow prepares specialized metabolite spectra for visualization in the interactive analysis platform. Spectra are smoothed with the Savitzky Golay filter (half window size=20) and baseline corrected via the TopHat method, peaks were detected using the SuperSmoother algorithm with a half window size of 20 and a signal-to-noise threshold of 1.^52^ A media subtraction is applied, removing all peaks in spectra that are within 0.001 Da of peaks that appear across all scans in the control spectra. Finally, intensities are summed across replicates and normalized with a base peak set to 1.0. Finally, peaks are discretized into 1 Da bins to account for limitations in instrument mass accuracy (see **Supplementary Note 3**).

### Strain deposition Workflow

The IDBac deposition workflow enables users to validate and deposit spectra from known strains into the IDBac-KB. During validation, the spectrum processing is mirrored from the knowledgebase search. Once validated, users can disable the dry-run mode, allowing the raw spectra to be sent to IDBac-KB as JSON objects containing peak lists and metadata (e.g., 16S rRNA gene sequence, Genbank accession numbers, taxonomy IDs, and preparation conditions) where they are associated with a unique identifier.

### IDBac-Knowledgebase

The internal knowledgebase contains two datasets: raw spectra and processed spectra. When new spectra are deposited, a data transformation and enrichment workflow integrates taxonomic information from Genbank and NCBI taxonomy databases into the JSON objects and generates a metadata table viewable in the database portal. Additionally, this workflow processes each deposited strain in an identical manner to the analysis workflow, mitigating duplicated computation between analysis workflow queries.

The interactive knowledgebase portal allows users to explore the diversity of deposited spectra and associated metadata. A continually updated taxonomic tree is recreated using the ETE 3 toolkit^64^ from associated NCBI taxonomy IDs. The interactive interface is implemented in Python using the Dash and Plotly packages.^65^

### Interactive Analysis Platform

The interactive analysis platform allows users to explore the results of their protein spectra search results along with their associated metabolites through a Streamlit app. The interactive protein dendrogram, implemented with Plotly and SciKit Learn^66^ Python packages, allows users to explore different approaches toward hierarchical clustering of protein isolates (e.g., average, single, and complete linkage). Specifically, the workflow applies the user-specified pairwise distance function to generate clustering analysis based on processed consensus spectra (see **Methods: IDBac-Analysis Workflow**). Additionally, it allows users to specify the number of database hits by both distance and number, allowing for a focused analysis of top candidates. Finally, user-uploaded metadata can be displayed alongside the dendrogram as both text and scatter plots enabling the study of the interplay between experimental conditions and spectral similarity.

The protein heatmap page allows users to select and plot large numbers of strains side-by-side. Strains can be selected individually or based on metadata and clustering labels. The displayed *m/z* features can be filtered by a variety of criteria including relative intensity, count between strains, and replicate presence. The replicate presence threshold can be modified and allows users to specify either a ppm or fixed tolerance for counting replicates in neighboring bins. Notably, tolerances are applied to binned spectra, meaning the lower bound for tolerance must be less than the minimum bin *m/z* feature, and the upper bound higher than the maximum bin *m/z* feature.

Similarly, the metabolite association network page allows for the interactive exploration of metabolites associated with analyzed strains both through a heatmap and MAN visualization. The displayed specialized metabolites can be filtered by *m/z*, relative intensity, and replicate frequency. The visualization can additionally be customized by node color and shape based on protein cluster and metadata values.

### Isolation and Identification of *Paraburkholderia acidicola* RL17-338-BIF-B

*Paraburkholderia acidicola* RL17-338-BIF-B (1-D8) was isolated from a rhizome associated with plant roots collected near Soames Hill, Gibsons, British Columbia, Canada (latitude: 49.42° N; longitude: 123.49° W; altitude: 172 m). The environmental sample was plated onto BIF isolation agar (15 g/L agar, 3 g/L monopotassium phosphate, 0.2 g/L magnesium sulfate heptahydrate, 5 g/L L-threonine, 100 mg/L fusaric acid, 100 mg/L bacitracin, 100 mg/L cycloheximide, 100 mg/L nystatin, 100 mg/L econazole, 0.292 g/L copper (II) sulfate, 0.085 g/L nickel (II) sulfate heptahydrate) and individual colonies isolated on LB media. Frozen stocks of overnight liquid culture were created in 1:1 glycerol/water in cryomicrocentrifuge tubes and stored at −70°C.^67^

Initially identified as *Paraburkholderia megapolitana* via 16S rRNA gene sequencing, RL17-338-BIF-B was reassigned to *P. acidicola* after the whole genome sequence of the strain was submitted to GenBank, available under the accession code JAOALG000000000 and BioProject ID number PRJNA875462.^68^

### Cultivation and Extraction

*P. acidicola* RL17-338-BIF-B was cultured from frozen stocks on nutrient agar (NA) plates for 2 days at 30 °C and subsequently inoculated into 10 mL BYP-glucose medium (5 g/L starch, 4 g/L yeast extract, 2 g/L peptone, 3 g/L sodium chloride, 5 g/L D-glucose) in 40 mL culture tubes. This seed culture was incubated in an incubator-shaker overnight (200 rpm, 27 °C). 3 mL of seed culture was then inoculated into 57 mL BYP-glucose in 250 mL wide-neck Erlenmeyer flasks with metal springs covered with milk filters and incubated for three days on a rotary shaker (200 rpm, room temperature). A final fermentation period was started by inoculating 45 mL of medium-scale culture into 2.8 L Fernbach flasks containing large metal springs and 1 L BYP-glucose supplemented with 20 g Amberlite XAD-16 adsorbent resin. This large-scale culture was fermented for five days (200 rpm, room temperature), after which the resin was collected by vacuum filtration using glass microfiber filters. 1:1 DCM/MeOH was added to the vacuum-dried filters and resin and the mixture was filtered under vacuum. The resulting filtrate was dried by rotary evaporation, then resuspended in MeOH and combined with 10 g Celite for each liter of culture extracted. The mixture was dried by rotary evaporation and the dried Celite loaded into a cartridge for separation with a 5 g C18 column on a CombiFlash Rf system (Teledyne Isco) using a MeOH/H_2_O step-gradient method to collect seven prefractions: 10% MeOH, 20% MeOH, 40% MeOH, 60% MeOH, 80% MeOH, 100% MeOH, and 100% EtOAc.

### Purification of Haereoacidicolin

Haereoacidicolin was purified from the 60% and 80% MeOH prefractions via reversed-phase HPLC on an Agilent Technologies 1200 series HPLC system using the following conditions: Kinetex 5 µ XB-C18 100 Å (250 × 4.6 mm); 1 mL/min flow rate; isocratic 21% HPLC-grade MeCN (Sigma-Aldrich) + 0.02% formic acid mobile phase; UV detection at 254 nm.

### Acquisition of Spectra

The UV-Vis spectrum was obtained on an Evolution™ 260 Bio UV-Visible Spectrophotometer (Thermo Fisher). Optical rotation was measured on a Model 341 Polarimeter (PerkinElmer). HR-ESI-MS and MS2 fragmentation spectra were obtained on a SYNAPT UPLC-ESI-qTOF (Waters Corporation). NMR data were acquired at 25 °C using a Bruker Avance III 600 MHz spectrometer equipped with a 5 mm QCI cryoprobe. Spectra were referenced to residual non-deuterated DMSO (δ_H_ 2.50/δ_C_ 39.5).

### Marfey’s Analysis of Haereoacidicolin

0.2 mg of haereoacidicolin was hydrolyzed with 6N HCl at 110 °C for 18 hours. The solution was concentrated to dryness under a stream of N_2_ gas. The hydrolysate was re-suspended in 100 µL of 50% aqueous acetone and derivatized with 100 µL of Nα-(2,4-dinitro-5-fluorophenyl)-L-valinamide (L-FDVA, 10 mg/mL acetone, Sigma-Aldrich) under basic conditions (40 µL of 1M NaHCO_3_, Fisher Chemical) at 40 °C for 1 hour. The reaction mixture was cooled to room temperature and neutralized with 1N HCl (40 µL), then concentrated to dryness under a stream of N_2_ gas. The derivatized haereoacidicolin (**1**) was re-suspended in MeOH (100 µL, Fisher Chemical Optima™ LC/MS-grade) for analysis via UPLC-MS (RDa, Waters) using an ACQUITY UPLC® and HSS T3 column (2.1 × 100 mm, 1.8 µm, 0.5 mL/min) with a gradient solvent system of 5% MeCN (with 0.01% formic acid) to 98% MeCN (with 0.01% formic acid) over 13 min at a flow of 0.5 mL/min. The following amino acid standards were derivatized, prepared and analyzed in the same manner as the haereoacidicolin (**1**) hydrolysates mixture: D-Ser (Sigma-Aldrich), L-Ser (ACROS ORGANICS), D-Glu (Sigma-Aldrich), L-Glu (Tokyo Chemical Industry Co., Ltd), D-Thr (Alfa Aesar), L-Thr (Sigma-Aldrich), D-*allo*-Thr (Alfa Aesar), and L-*allo*-Thr (Sigma-Aldrich). The amino acids in the structure were assigned through analysis using Waters UNIFI software.

### Characterization of Haereoacidicolin

Haereoacidicolin was isolated as yellow solid; [α]_D_^20^ +8.7 (c 0.26, MeOH); UV (MeOH) λmax (log ε) 270 nm (2.67); ^1^H and ^13^C NMR data (600 MHz and 150 MHz, DMSO-d_6_), see **Supplementary Table S4**; HR-ESI-MS [M + H]^+^ *m/z* 764.3858 (calculated for C_35_H_54_N_7_O_12_^+^, 764.3825). All raw NMR data files were deposited in the open access Natural Products Magnetic Resonance Database (www.np-mrd.org).^69^

## Data Availability and Materials

### Documentation

https://sites.google.com/uic.edu/idbac-documentation/

### Source Code

Source code is available in the following repositories: IDBac Analysis Interface (https://github.com/Wang-Bioinformatics-Lab/IDBac_Interactive_Interface), IDBac Deposition Workflow (https://github.com/Wang-Bioinformatics-Lab/IDBacDeposition_Workflow), IDBac Analysis Workflow (https://github.com/Wang-Bioinformatics-Lab/IDBac_Analysis_Workflow), IDBac KB Server (https://github.com/Wang-Bioinformatics-Lab/IDBac-KB-Server). Additionally, source data are provided with this paper.

## Author contributions

NK, MS, LMS, MW and BTM were responsible for platform development. NK and BTM were responsible for data analysis. MS and MW developed the computational methods, wrote software, conducted statistical analyses, and contributed to data analysis. NK was responsible for coordinating depositions of the initial knowledgebase population. NK, MT, GM, and JR prepared MALDI-TOF MS target plates for data collection and compiled metadata for knowledgebase depositions. NK acquired MALDI-TOF MS protein data for knowledgbase depositions. NK, SR, AH, and BTM were responsible for MALDI-TOF analysis on *Paraburkholderia* spp. JS, LZ, CF, and RL were responsible for the isolation and structure elucidation of haereoacidicolin. BP, SR, ASE were responsible for BGC analysis of haereoacidicolin. HB and DC provided samples that were used for analyses in Fig. 4. NK, RS, DB, SP, JR, AFS, GM, AB, CF, CK, MWM, CD, AB, CC, LG, CS, JMW, BIA, WM, EGP TB, MH, RL, LS, and BTM contributed strains to the IDBac-KB. NK, MS, BTM, MW, and LMS wrote the manuscript.

## Acknowledgements

This work was supported by the Chicago Biomedical Consortium with support from the Searle Funds at The Chicago Community Trust (BTM and Neil Kelleher); by the National Institute of General Medical Sciences of the NIH award R01GM125943 (BTM and LMS), R21GM148870 (LMS), R01GM129344 (ASE, BTM and RGL), R25GM051765 (CDS), R01AI155694 (JMW), R35GM147235 (SMKM), SC2GM144172 (EPM) and by National Science Foundation graduate research fellowship program (RAS). MS, MW was supported by NIH 5U24DK133658-02. SR and GM were supported by the Office of the Director and the National Center for Complementary and Integrative Health under T32AT007533. MS, MW was supported by NIH 5U24DK133658-02. MW was partially supported by the U.S. Department of Energy Joint Genome Institute (https://ror.org/04xm1d337), a DOE Office of Science User Facility, is supported by the Office of Science of the U.S. Department of Energy operated under Contract No. DE-AC02-05CH11231. NG and MML were supported by NIH R35GM150870 to NG. The content is solely the responsibility of the authors and does not necessarily represent the official views of the National Institutes of Health. The authors wish to acknowledge the Integrated Molecular Structure Education and Research Center (IMSERC) at Northwestern University where MALDI-TOF MS for database deposition data were collected. The authors wish to acknowledge Oscar Spilberg for collecting moss and cockroach samples; Emilie Blevins, M.S. for collecting freshwater sponge samples; and Dr. Maxim Svetlov for collecting moss and forest detritus samples. Further, the authors wish to acknowledge the following researchers for knowledgebase contributions: Dr. Valerie Paul, Dr. Bonnie K. Baxter, and Dr. Vinayak Agarwal. Finally, the authors wish to acknowledge Dr. Chase Clark for his introduction of the original IDBac desktop platform and support in developing the IDBac-KB and web-based analysis platform.

