## Supplementary Information for "IDBac: an open-access web platform to identify bacteria and analyze relationships in culture collections using MALDI-TOF mass spectrometry"

[Figure S14. 1D selective ROESY NMR spectrum, irradiating 12-NH (δ_H_ 8.28), of haereoacidicolin in
 DMSO-d_6_ at 600 MHz.	29](#_Toc222139509)

[Figure S15. 1D selective ROESY NMR spectrum, irradiating 8-NH (δ_H_ 9.76), of haereoacidicolin in
DMSO-d_6_ at 600 MHz.	30](#_Toc222139510)

### **Table S1.** **A comparison of several currently available analysis platforms and spectral repositories.**

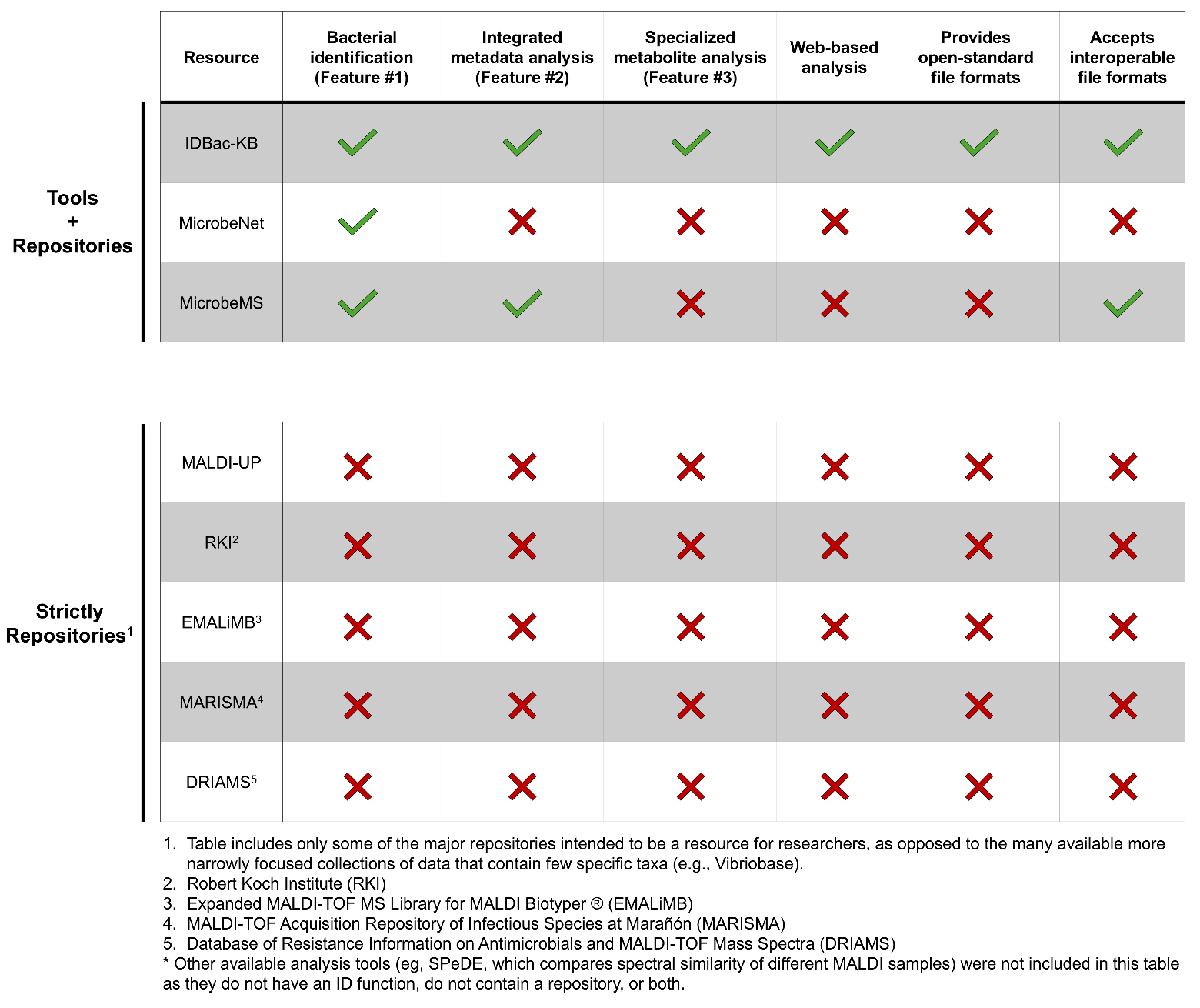

**A comparison of several currently available analysis platforms and spectral repositories.** Existing large-scale MALDI-TOF data repositories typically consist of vendor based proprietary formats primarily including .btmsp and fid. In contrast, IDBac-KB uses the open-standard .mzML file format that enables broad interoperability between instrument vendors. Analysis platforms that exist behind a paywall (e.g., Biotyper) are not included here.

### **Figure S1. Search results for *Solwaraspora* sp. Z001 query strain.**

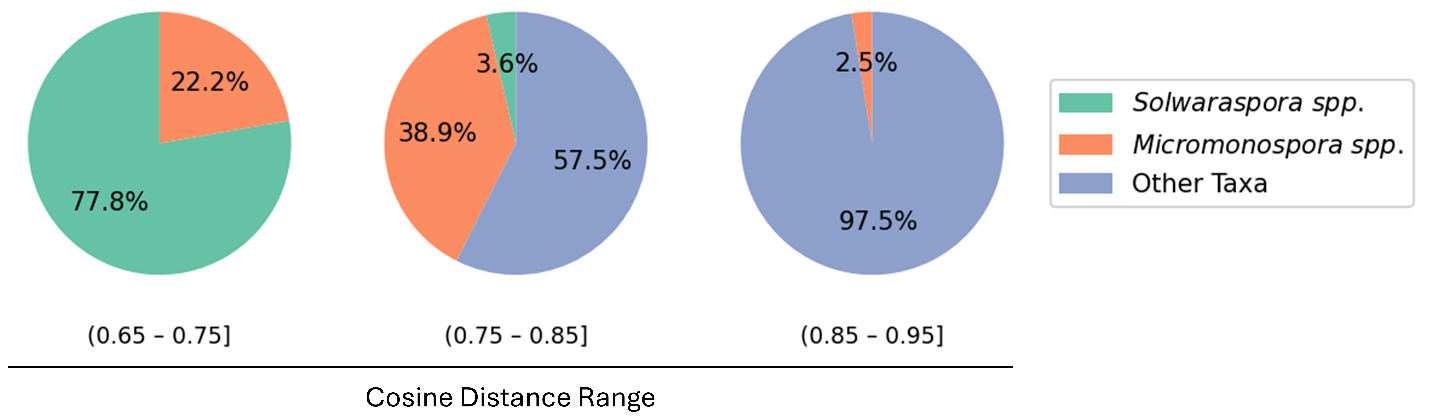

Pie charts summarize IDBac query results for isolate Z001 (identified as a *Solwaraspora* sp*.*) at various search thresholds; see main text **Fig 3A**. At a threshold of ≤0.75, the query strain matched exclusively with *Solwaraspora* sp*.* and *Micromonospora* sp*.* KB entries. We note that members of the family; *Micromonosporaceae* possess a high degree of genetic homology and genus level distinction is an ongoing debate amongst the community.^1^ For example, The Genome Taxonomy Database, currently classifies *Solwaraspora* sp*.* as *Micromonospora_E* sp*.*, while the NCBI Taxonomy Database and List of Prokaryotic names with Standing in Nomenclature distinguishes between the two genera. Emerging evidence suggests that *Micromonospora_E* strains should be reclassified as an independent genus,^2^ though the field has not yet reached a consensus.

### **Table S2. Cosine distances between query isolates and IDBac-KB strains.**

Isolate accession numbers and associated metadata can be found at <https://idbac.org/knowledgebase>.

| **Blind Query ID** | **Query ID** | **Query taxonomy** | **IDBac-KB ID** | **IDBac-KB taxonomy** | **Cosine distance** |
| --- | --- | --- | --- | --- | --- |
| ^▲^C | MTH1 - G07 | *Bifidobacterium longum* | ^●^B-5799 | *Streptomyces phaeoluteichromatogenes* | 0.6880 |
| ^▲^D | MTH1 - G08 | *Bifidobacterium longum* | ^*^MSK.5.11 | *Bifidobacterium longum* | 0.68278 |
| ^▲^E | MTH2 – A08 | *Bacteroides thetaiotaomicron* | *MSK.23.25 | *Bacteroides thetaiotaomicron* | 0.6085 |
| ^▲^F | MTH2 – C01 | *Bacteroides thetaiotaomicron* | *MSK.23.25 | *Bacteroides thetaiotaomicron* | 0.6044 |
| ^▲^G | MTH2 - C02 | *Bacteroides caccae* | *MSK.18.53 | *Bacteroides caccae* | 0.6102 |
| ^▲^H | MTH2 - C07 | *Bacteroides eggerthii* | *MSK.16.19 | *Bacteroides eggerthii* | 0.5339 |
| ^▲^I | MTH2 - B03 | *Bacteroides xylanisolvens* | *MSK.19.24 | *Bacteroides xylanisolvens* | 0.6610 |
| ^▲^J | MTH2 - D02 | *Bacteroides fragilis* | *MSK.7.6 | *Bacteroides fragilis* | 0.6094 |
| ^▲^K | MTH1 - D03 | *Eubacterium contortum* | ^●^ISP-5022 | *Streptomyces filamentosus* | 0.7124 |
| ^▲^L | MTH2-E03 | *Clostridium innocuum* | ^+^502A | *Staphylococcus aureus* | 0.7416 |
| ^▲^M | MTH1 - F06 | *Lactococcus lactis* | ^#^Cn52-H1 | *Vibrio coralliilyticus* | 0.7079 |
| ^▲^N | RJX 1119 | *Mediterraneibacter gnavus* | *MSK.15.77 | *Mediterraneibacter gnavus* | 0.6328 |
| ^▲^O | RJX 1120 | *Mediterraneibacter gnavus* | *MSK.5.17 | *Mediterraneibacter gnavus* | 0.6061 |
| ^▲^P | RJX 1123 | *Mediterraneibacter gnavus* | *MSK.5.17 | *Mediterraneibacter gnavus* | 0.6184 |
| ^▲^Q | RJX 1126 | *Mediterraneibacter gnavus* | *MSK.15.77 | *Mediterraneibacter gnavus* | 0.6145 |
| ^▲^R | MTH1 - B07 | *Blautia obeum* | ^$^MER 118 | *Bacillus cereus* | 0.7415 |
| ^▲^S | MTH1 - C05 | *Mediterraneibacter gnavus* | *MSK.23.82 | *Mediterraneibacter gnavus* | 0.5755 |
| ^▲^T | MTH2 - F04 | *Escherichia fergusonii* | *DFI.3.23 | *Phocaeicola vulgatus* | 0.6863 |

^▲^Strains obtained from the laboratory of Dr. Matthew Henke, University of Illinois Chicago.

^★^Strains obtained from the laboratory of Dr. Roger Linington, Simon Fraser University.

*Strains obtained from the Duchossois Family Institute at the University of Chicago.

^●^Cell material obtained through collaboration with the laboratory of Dr. William M. Metcalf, University of Illinois Urbana-Champaign and the U.S. Department of Agriculture.

^■^Strain obtained from the laboratory of Dr. Brian T. Murphy, University of Illinois Chicago.

^+^Spectra submitted by the laboratory of Dr. Neha Garg, Georgia Institute of Technology

^#^Strains obtained from the laboratory of Valerie Paul. Spectra submitted by the laboratory of Dr. Neha Garg, Georgia Institute of Technology

^$^Strains obtained through a collaboration with Dr. Adriana Blachowicz and the NASA Jet Propulsion Laboratory.

### **Figure S2. IDBac-KB genus-level discrimination of queried microbiome isolates.**

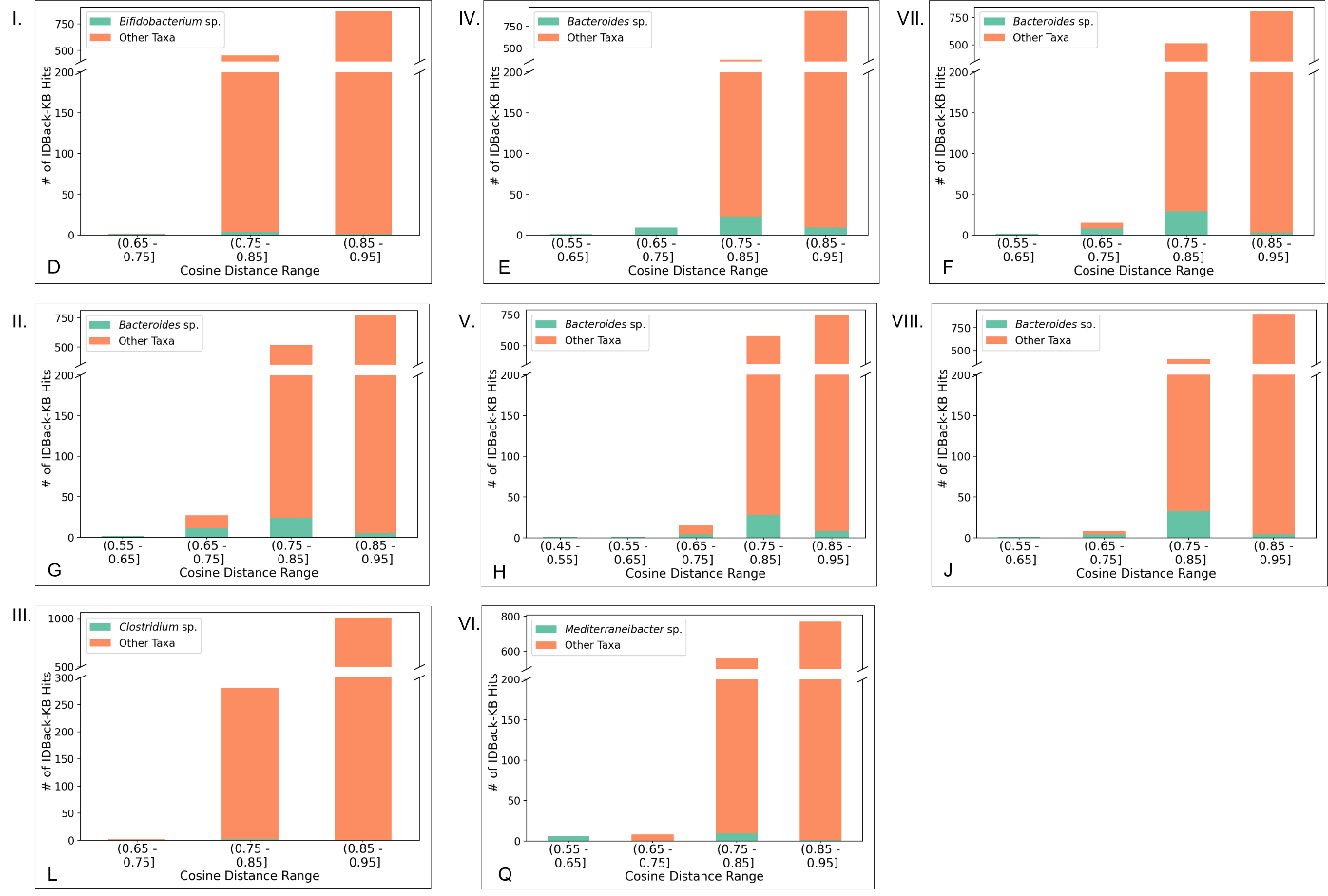

**Genus-level discrimination in KB search.** A stacked bar chart demonstrates the proportion of intra- (green) and inter-genus (orange) hits binned by intensity-agnostic cosine distances. Query isolate IDs are listed in the bottom left corner of each chart. Example: Hits produced for *Bifidobacterium* sp*.* query strain D (I.), hits produced for *Bacteroides* sp*.* query strain E (IV), etc. Lower cosine distances denote more similar spectral matches. Across all examples, matches under a distance of 0.65 maintain perfect precision. Higher thresholds include additional correct hits but come at the cost of lower precision. For example, for the 0.45-0.65 range, all hits are within-genus for a *Bacteroides sp.* query strain, H (V.). However, only 36.36% and 5.09% of hits are within-genus for the 0.65-0.75 and 0.75-0.85 bins, respectively.

### **Table S3. IDBac-KB Hits for microbiome data at various search thresholds.**

| **Strain ID** | **<0.65  Correct genus** | **<0.65 Different genus** | **0.65-0.70 Correct genus** | **0.65-0.70**  **Different genus** | **0.70-0.75  Correct genus** | **0.70.-0.875  Different genus** |
| --- | --- | --- | --- | --- | --- | --- |
| C | - | - | - | 2 | 2 | 40 |
| D | - | - | 1 | 0 | 0 | 1 |
| E | 1 | - | 5 | - | 4 | - |
| F | 2 | - | 7 | - | 1 | 7 |
| G | 2 | - | 3 | - | 8 | 16 |
| H | 2 | - | 2 | - | 2 | 11 |
| I | - | - | 4 | - | 4 | 1 |
| J | 1 | - | 2 | - | 2 | 4 |
| K | - | - | - | - | - | 4 |
| L | - | - | - | - | 1 | 1 |
| M | - | - | - | - | - | 16 |
| N | 3 | - | 3 | 1 | - | 13 |
| O | 2 | 1* | 3 | 1 | 1 | 6 |
| P | 4 | - | 2 | 1 | 3 | 2 |
| Q | 6 | - | - | 1 | - | 7 |
| R | - | - | - | - | - | 2 |
| S | 6 | - | - | - | - | 8 |
| T | - | - | - | 1 | - | 10 |

*Hit outside of target genus: MSK.22.44 | JAKNIV000000000 | Klebsiella pneumoniae*.*

**KB hits for microbiome data at various search thresholds.** KB search results for all microbiome strains queries (see main text **Fig. 3B**) using the intensity-agnostic cosine distance metric. In all but one case, hits below a distance threshold of 0.65 had 100% precision. Notably, the K and M query strains served as negative controls since no strains within-genus were contained in the KB for the query strains to match.

### **Figure S3. Example of relative intensity-driven protein heatmap.**

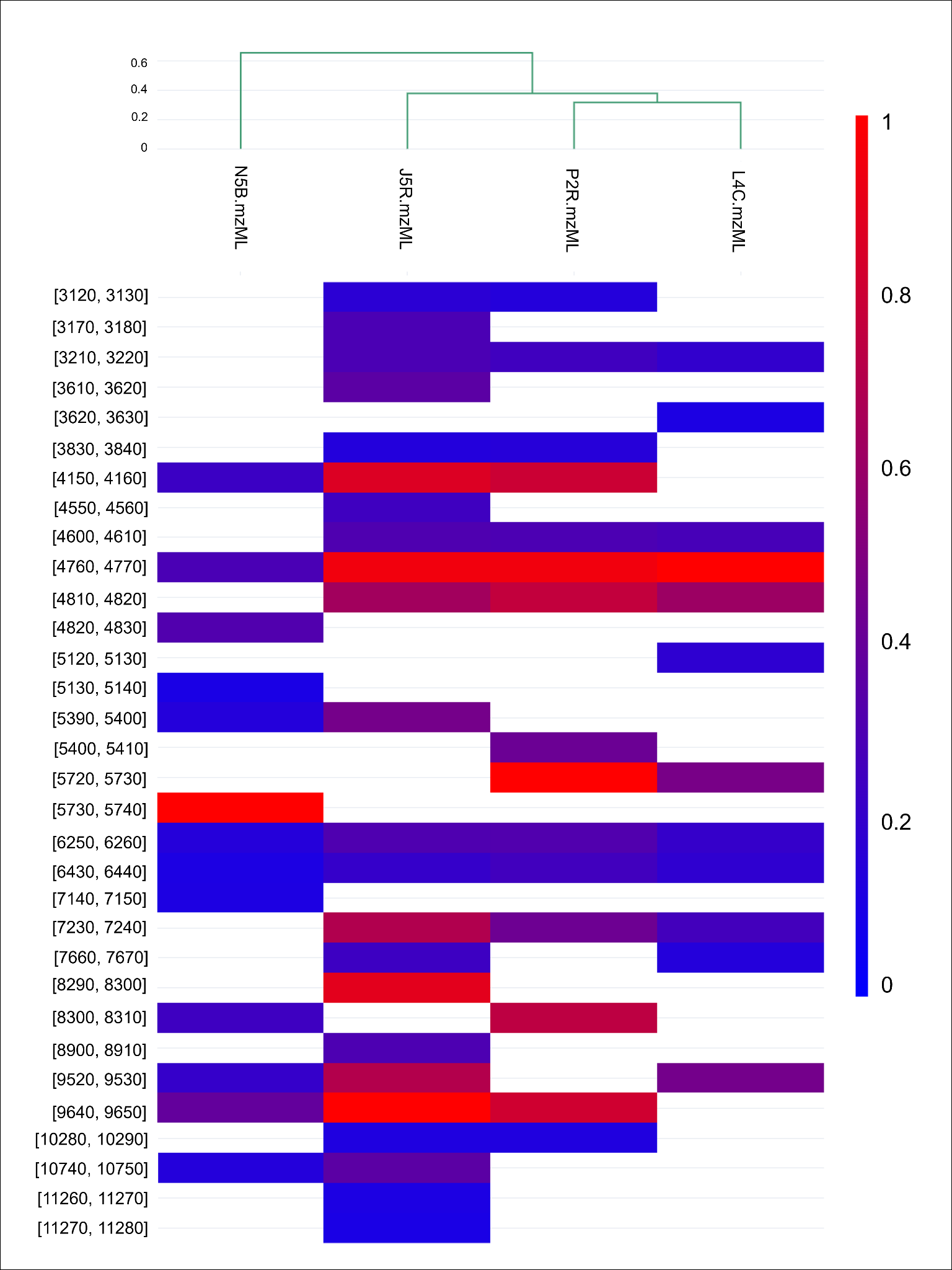

**Heatmap of Protein Peak Intensities.** A heatmap showing shared MALDI MS *m/z* values of isolates derived from four soil samples collected from the rhizosphere of plant samples in Maine. Intensities were thresholded at 50% presence in replicates and a relative intensity of 0.10. This revealed the *m/z* values that drove clustering of the isolates.

### **Supplementary Note 1A.**

Pairwise differences in MALDI spectral profiles between isolate sources (lake: n=56, sediment: n=23 samples) were assessed using a PERMANOVA on cosine distance matrices (pseudo-F=9.633738, p=0.001, n=79, 999 permutations). The result indicates that isolates from different sources exhibit significantly different MALDI spectral profiles, which could be due to a shift in the mean spectral features (centroid), a change in the variability of spectra (dispersion), or a combination of both. To distinguish these possibilities, a subsequent PERMDISP test for multivariate homogeneity of variances confirmed a significant difference in dispersion (variability) of the spectral profiles between the two isolate sources (F=10.55, p=0.001). Therefore, the significant PERMANOVA result is driven, at least in part, by differences in spectral heterogeneity (i.e., isolates from one source are more variable in their spectral profiles when compared with the other source). To further quantify the effect of isolate source on proteomics profiles, a Principal Coordinates Analysis (PCoA) was performed on the cosine distance matrix and the source centroids were calculated in the reduced space. The PCoA clearly illustrates the multivariate location shift with the centroids for the two sources separated along the primary axis (PC1) (See **1A** below).

Additionally, hierarchical clustering of cosine distances revealed strong concordance with source labels. Though notably, two samples were collected from each source, comprising four total sample locations on the left- and right-hand side of the lake (LHS, n=29; RHS n=27 samples), and in shallow (“Lake”, n=12 samples) and deeper water (“Lake – Deeper”, n=11 samples). To assess concordance with source location, singleton clusters at a cut height below 0.7 were removed from analysis and remaining spectra were reclustered using a max cluster criterion of four. This produced four independent clusters, each with a different dominant isolate source, and an average cluster purity of 0.68. When clustering was evaluated with a max cluster criterion of two and binary labels (Sediment or Lake Water), average cluster purity was 1.0, reflecting perfect agreement between cluster assignment and isolate source. To disambiguate whether clustering was a function of sample source or taxonomy, annotations provided by 16S sequencing were integrated. No common genera were found between lake and sediment sources. However, sample source locations were mixed within any given genus, suggesting that the observed separation may be a function of species-level variations.

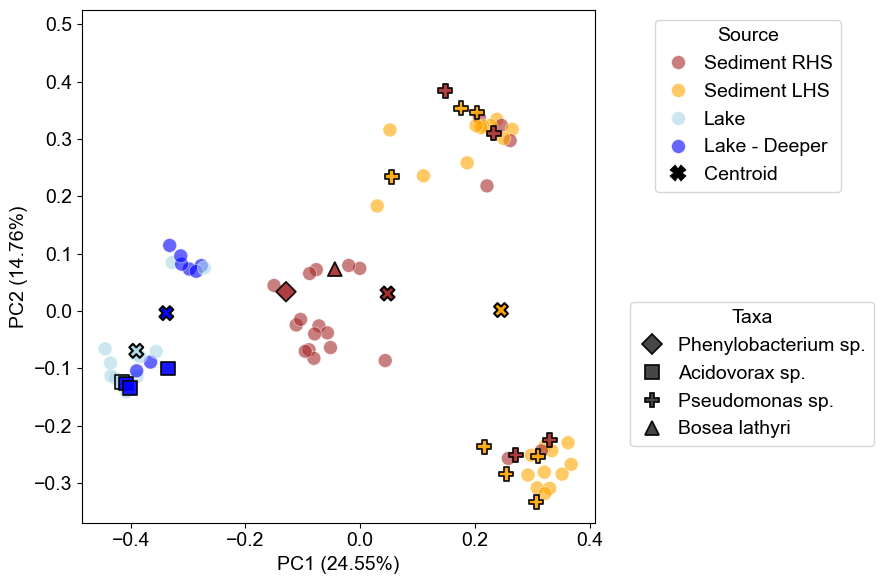

**1A.** PCoA plot shows the multivariate spectral profiles. Each point represents an individual isolate, colored by its source (Sediment or Lake water) and collection location. Markers denote 16S annotations. Separation between isolate sources (sediment/lake) can be observed along the first principal component. *Acidovorax* sp. were identified exclusively in the lake water, while all other sequenced strains were identified in sediment.

### **Supplementary Note 1B.**

Pairwise differences in multivariate proteomic profiles among the four isolate collection time points (6 hours: n=37; 30 hours: n=140; 78 hours: n=49; 102 hours: n=47 samples) were significant (PERMANOVA: pseudo-F = 8.47, p = 0.001, n = 273, 999 permutations). However, a subsequent test for homogeneity of multivariate dispersions also indicated significant differences in within-group variance (PERMDISP: F = 220.73, p = 0.001), suggesting that part of the observed separation may be attributable to differences in dispersion among sources (see Main Text Fig. 4C). To quantify the temporal effect, PCoA was performed on the cosine distance matrix and distance between centroids was calculated in the reduced space. The first three principal components (PC1, PC2, and PC3) collectively explain only 19.08% of the variance. Therefore, the PCoA plot serves as an illustrative projection of the differences, but it only captures a small fraction of the true dissimilarity. Nevertheless, the PCoA does illustrate a multivariate location shift primarily along the first principal component (PC1). Notably, across time points, PC1 becomes increasingly negative, with the largest shift between 30 hours observed between 30 and 78 hours. Again, hierarchical clustering was performed and revealed the relationship between dendrogram distances and isolation time. We report the proportion of clusters with purity=1.0 and the average cluster size as a function of the number of clusters in **1B**. Notably, as the number of clusters decreases from 274 (number of samples) to 100, the proportion of pure clusters remains high, at 83.17%, reflecting the concordance between dendrogram near-neighbors and isolation time.

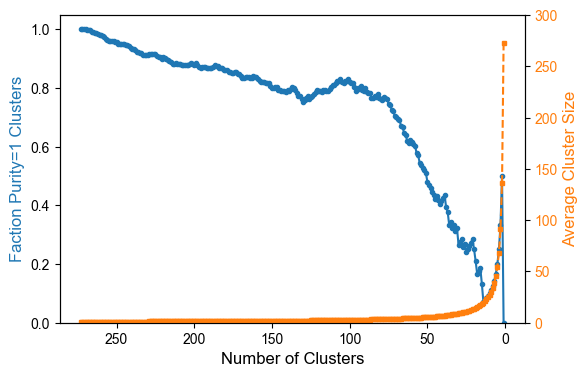

**1B**. Number of clusters vs fraction of perfectly pure clusters and mean cluster size. The tightest clusters (merged first) show a high rate of perfect purity examples, highlighting that dendrogram near-neighbors generally come from the same isolation time point.

### **Supplementary Note 2.**

Nodes in metabolite association networks (MANs) represent *m/z* values associated with strains, and since MALDI is a ‘soft’ ionization technique, these values often represent intact masses and singly charged adducts such as the protonated molecule, [M+Na]^+^, [M+K]^+^. In some instances, nodes may represent shared metabolites, however it is also possible that they represent isobaric compounds. In this instance further resolution may be obtained by high resolution tandem mass spectrometry (HRMS/MS) or using further separation such as ion mobility spectrometry (IMS).

Subtraction of a matrix blank is employed to remove *m/z* signals attributable to the matrix, however we acknowledge that select few metabolite signals may be filtered out in this process. Several studies from our team indicate that this does not significantly skew the interpretation of MANs ([Shepherd et al, Anal Chem, 2025](https://pubs.acs.org/doi/10.1021/acs.analchem.5c02787); [Luu et al, Analyst, 2023](https://pubs.rsc.org/en/content/articlelanding/2023/an/d3an00408b); [Clark et al, ISME Comm 2022](https://www.nature.com/articles/s43705-022-00105-8); [Clark et al, Molecules 2022](https://www.mdpi.com/1420-3049/27/7/2038); [Condren et al, Gut Microbes, 2020](https://www.tandfonline.com/doi/full/10.1080/19490976.2020.1740073); [Costa et al, J Nat Prod, 2019](https://pubs.acs.org/doi/10.1021/acs.jnatprod.9b00168); [Clark et al, Proc Natl Acad Sci, 2018](https://www.pnas.org/doi/10.1073/pnas.1801247115)).

### **Figure S4. MS/MS similarity of haereomegapolitanin A and 3-E9.**

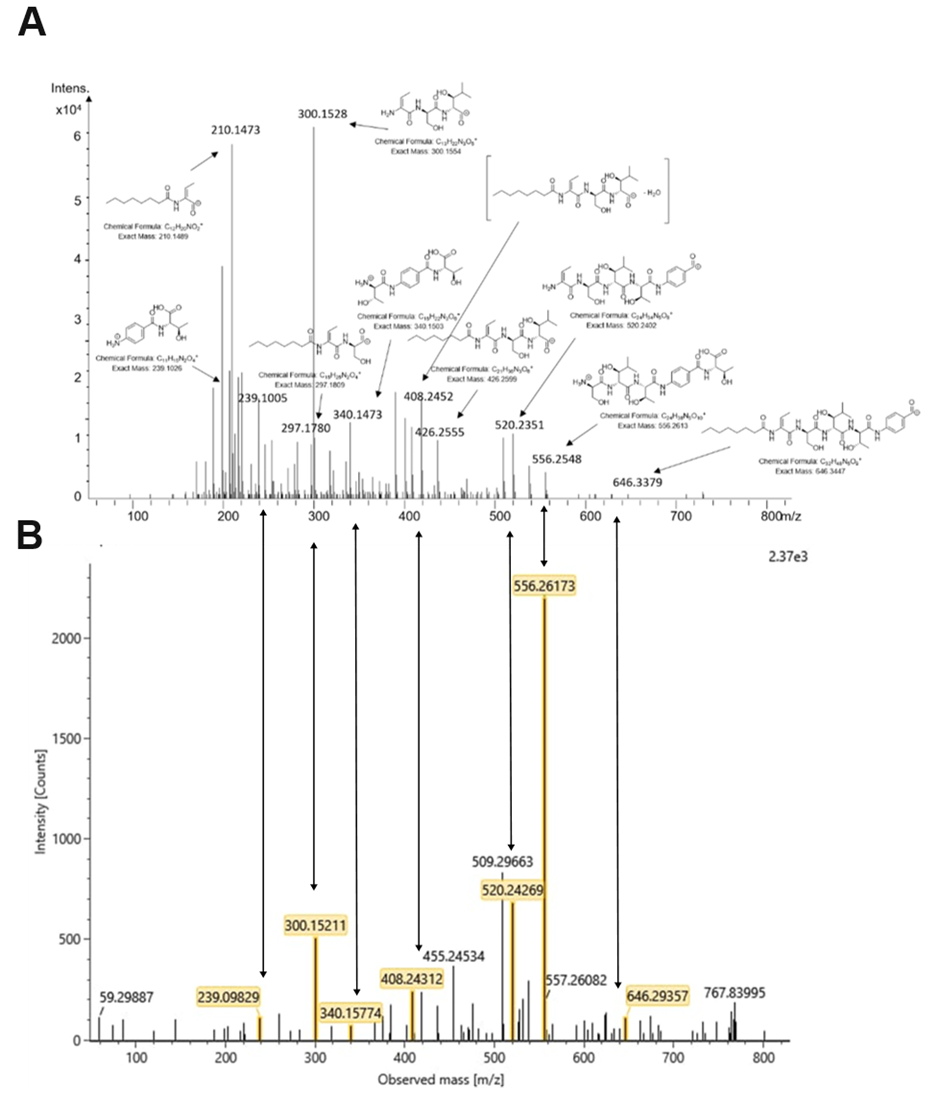

**A)** MS/MS fragmentation of haereomegapolitanin A from original discovery by Zheng et al.^3^ Reprinted with permission under the terms of the Creative Commons Attribution-Non Commercial License. **B)** MS/MS fragmentation of haereomegapolitanin A produced by strain 3-E9. Double-headed arrows point toward matching fragments.

### **Figure S5. Haereoacidicolin MS/MS fragmentation.**

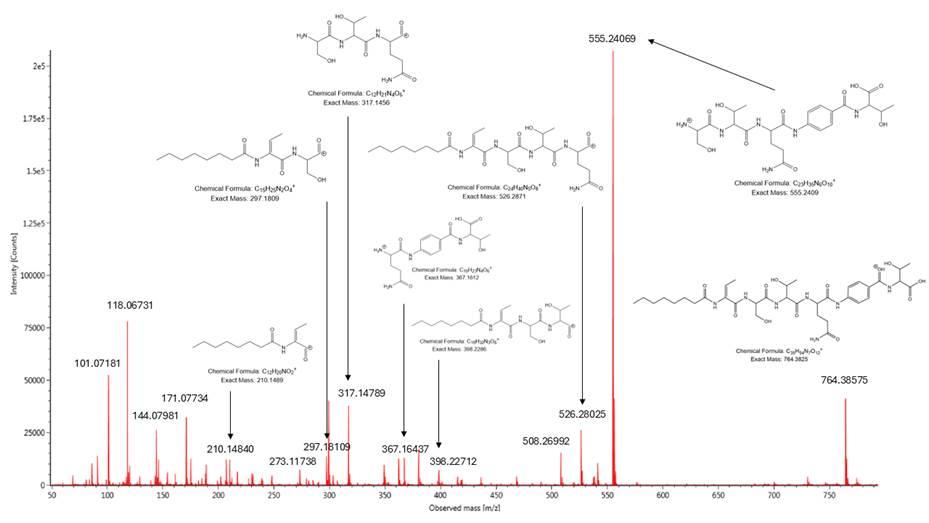

### **Supplementary Note 3.**

**Planar structure elucidation of haereoacidicolin from NMR data.**

The molecular formula was determined as C_35_H_53_N_7_O_12_ by HR-ESI-MS ([M+H]^+^ *m/z* 764.3858) and NMR (Table S3), indicating 13 degrees of unsaturation. The ^1^H and phase-sensitive ^1^H-^13^C HSQC spectra revealed 4 × CH_3_, 9 × CH_2_, and 11 × CH, which includes signals for two pairs of equivalent aryl protons indicating a para-substituted aromatic ring. In addition, the ^1^H and ^1^H-^15^N HSQC spectra revealed 1 × primary amide NH_2_ and 6 × secondary amide NH signals. Evaluation of the ^13^C and HMBC spectra further identified 11 × qC, including one olefinic carbon, two aryl carbons, and eight carbonyls. The remaining four hydrogen signals were attributed to hydroxy groups.

To solve the planar structure, the COSY and HMBC spectra were used to identify spin systems. HMBC correlations connected amide protons—8-NH (δH 9.76), 12-NH (δH 8.28), 15-NH (δH 8.04), 19-NH (δH 7.97), 23-NH2 (δH 6.77, 7.19), 24-NH (δH 9.85), and 29-NH (δH 7.81)—to their corresponding carbonyls and served as reference points for elucidating amino acid constituents within the molecule. Sequential COSY and HMBC correlations from methyl H3-1 (δH 0.81) to deshielded methylene H2-7 (δH 2.18) revealed an octanoate (Oct) adjacent to amide carbonyl C-8 (δC 172.1). An HMBC correlation from amide proton 8-NH to vinylic carbon C-9 (δC 131.8) started a spin system composed of the C-9 to C-11 positions, and correlations from vinylic proton H-10 (δH 5.56) to C-9 and methyl protons H3-11 (δH 1.76) revealed the dehydrobutyrine (Dhb) moiety which, combined with Oct, is characteristic of haereo compounds. Analysis of the 2D NMR data identified the presence and placement of amino acid residues Ser/Thr/Gln in order succeeding the Oct/Dhb. From the Gln α-proton H-20 (δH 4.30), an HMBC correlation to carbonyl C-24 (δC 170.4) connected the Gln residue to the spin system beginning with 24-NH (δH 9.85). Sequential HMBC correlations from 24-NH up to C-29 (δC 165.5) constructed the para-aminobenzoate (PABA), also characteristic of haereo compounds. Finally, COSY correlations from 29-NH (δH 7.81) to H3-32 (δH 1.06) and the deshielded chemical shifts of the methines H-30 (δH 4.24) and H-31 (δH 4.12) suggested the terminal Thr, however, the hydroxy proton signals and connections to a carboxylic acid carbonyl were not evident. Instead, the presence of a second Thr was corroborated by MS/MS measurements and Marfey’s analysis. A summary of key NMR correlations is presented in Figure S6.

**Configurational analysis of haereoacidicolin.**

The configuration of Dhb was determined using ROESY NMR (Figure S14 to Figure S15). Crosspeaks between amide proton 8-NH (δ_H_ 9.76) and vinylic proton H-10 (δ_H_ 5.56) and between the adjacent amide proton 12-NH (δ_H_ 8.28) and methyl protons H_3_-11 (δ_H_ 1.76) were observed, thus the configuration of the double bond was assigned as *E*.

The six stereogenic centers of haereoacidicolin were determined by Marfey’s method supplemented with analysis of the BGC (Figure S16-19, Figure 5B). The acid hydrolysate of haereoacidicolin and standards of the constituent amino acids-with glutamic acid in place of glutamine to account for its conversion during the hydrolysis of haereoacidicolin-were derivatized with Nα-(2,4-dinitro-5-fluorophenyl)-ʟ-valinamide (ʟ-FDVA). Comparison of LC/MS retention times indicated the presence of ᴅ-Ser, ʟ-Gln, ʟ-Thr, and ᴅ-*allo*-Thr residues. To ascertain the positions of each Thr residue, we analyzed the haereoacidicolin biosynthetic gene cluster using antiSMASH, and NaPDoS2^4^ with a focus on the substrate specificity of the adenylation (A) domains and the type of condensation (C) domains within the single nonribosomal peptide synthetase (NRPS) gene predicted to encode the peptide core present on haereoacidicolin. The predicted amino acids selected by the A domains were (N-terminus to C-terminus order, A1 to A5) l-Thr, l-Ser, l-Thr, l-Gln, and ‘unknown’, which was later assigned to a para-aminobenzoic acid moiety (PABA). The C-domains (Fig. S19) following each A domain led us to predict Dhb (mod C2 domain dehydrates l-Thr), d-Ser (dual C3 domain epimerizes l-Ser), d-Thr (dual C4 domain epimerizes l-Thr), and l-Gln (^L^C_L_ C5 domain). It is not currently possible to predict *allo*-Thr versus Thr incorporation, but Marfey's analysis clearly indicated d-*allo*-Thr, which we assigned to position A3. The l-Thr observed by Marfey’s analysis was then assigned to the Thr that offloads the peptide from the thioesterase domain and is attached to PABA. This offloading Thr cannot be predicted from the NRPS sequence.

### **Table S4. ^1^H (600 MHz) and ^13^C (150 MHz) NMR data for haereoacidicolin (1) in DMSO-d_6_.**

|  | **Position** | **δ_c_, type** | **δ_H_ (*J* in Hz)** | **HMBC** | **COSY** |
| --- | --- | --- | --- | --- | --- |
| **Oct** | 1 | 13.9, CH_3_ | 0.81, t (7.0, 7.0) | 2, 3 | 2 |
|  | 2 | 22.0, CH_2_ | 1.19, m^a^ | 1, 3, 4 | 1 |
|  | 3 | 31.1, CH_2_ | 1.15, m^a^ | 1, 2, 4, 5 |  |
|  | 4 | 28.6, CH_2_ | 1.23, m^a^ | 3, 5 | 6 |
|  | 5 | 28.4, CH_2_ | 1.16, m^a^ | 3, 4, 6, 7 | 6 |
|  | 6 | 24.9, CH_2_ | 1.46, m^a^ | 5, 7, 8 | 4, 5, 7 |
|  | 7 | 35.1, CH_2_ | 2.18, m^a^ | 5, 6, 8 | 6 |
|  | 8 | 172.1, C | – |  |  |
|  | 8-NH | – | 9.76, s | 8, 9, 10 |  |
| **Dhb** | 9 | 131.8, C | – |  |  |
|  | 10 | 118.0, CH | 5.56, q (7.3, 7.3, 7.4) | 9, 11, 12 | 11 |
|  | 11 | 12.8, CH_3_ | 1.76, d (7.3) | 9, 10, 12 | 10 |
|  | 12 | 164.9, C | – |  |  |
|  | 12-NH | – | 8.28, d (7.4) | 12, 13, 14 | 13 |
| **Ser** | 13 | 56.0, CH | 4.36, td (7.0, 7.0, 4.6) | 12, 14, 15 | 12-NH, 14 |
|  | 14 | 61.0, CH_2_ | 3.72, m |  | 13, 14-OH |
|  | 14-OH | – | 4.93, brd |  | 14 |
|  | 15 | 170.1, C | – |  |  |
|  | 15-NH | – | 8.04, d (7.4) | 15, 16, 17 | 16 |
| **Thr1** | 16 | 59.9, CH | 4.14, dd (6.0, 7.4) | 15, 17, 18, 19 | 15-NH, 17 |
|  | 17 | 66.8, CH | 3.98, ddq (4.3, 6.0, 7.0) | 15, 19 | 16, 17-OH, 18 |
|  | 17-OH | – | 5.26, d (4.3) |  | 17 |
|  | 18 | 20.0, CH_3_ | 1.16, m^a^ | 16, 17 | 17 |
|  | 19 | 170.2, C | – |  |  |
|  | 19-NH | – | 7.97, d (7.8) | 19, 20, 21 | 20 |
| **Gln** | 20 | 53.3, CH | 4.30, ddd (4.1, 7.6, 9.8) | 19, 21, 22 | 19-NH, 21a, 21b |
|  | 21a | 27.4, CH_2_ | 2.07, m^a^ | 20, 22, 23 | 22a |
|  | 21b |  | 1.76, m^a^ | 19, 20, 22, 23, 24 | 20, 22b |
|  | 22a | 31.6, CH_2_ | 2.18, m^a^ | 20, 21, 23 | 21a |
|  | 22b |  | 2.06, m^a^ | 20, 22, 23 | 21b |
|  | 23 | 173.6, C | – |  |  |
|  | 23-NH_2_ | – | 6.77, s | 22, 23 |  |
|  |  |  | 7.19, s | 23 |  |
|  | 24 | 170.4, C | – |  |  |
|  | 24-NH | – | 9.85, s | 24, 26 |  |
| **PABA** | 25 | 141.4, C | – |  |  |
|  | 26 | 118.6, CH | 7.73, d (8.4) | 25, 26, 28 |  |
|  | 27 | 127.9, CH | 7.82, d (8.4) | 25, 27, 29 |  |
|  | 28 | 129.0, C | – |  |  |
|  | 29 | 165.5, C | – |  |  |
|  | 29-NH | – | 7.81, brd | 29 | 30 |
| **Thr2** | 30 | 58.0, CH | 4.24, brd |  | 29-NH, 31 |
|  | 31 | 66.3, CH | 4.12, brd | 32, 33 | 32 |
|  | 31-OH | – |  |  |  |
|  | 32 | 20.0, CH_3_ | 1.06, m^a^ |  | 31 |
|  | 33 | 170.4, C | – |  |  |
|  | 33-OH | – |  |  |  |

^a^ Overlapped signals.

### **Figure S6. Key spectroscopic correlations of haereoacidicolin.**

**A)**
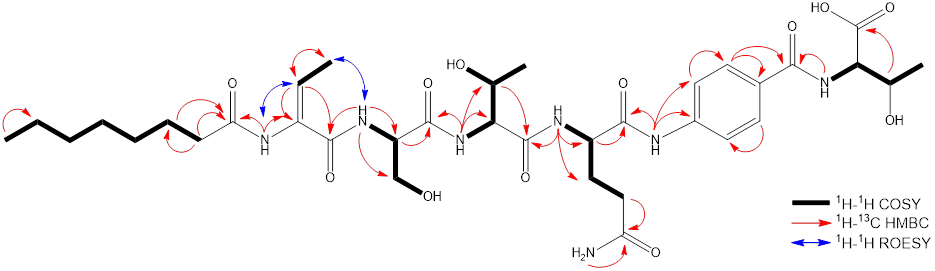

**B)**
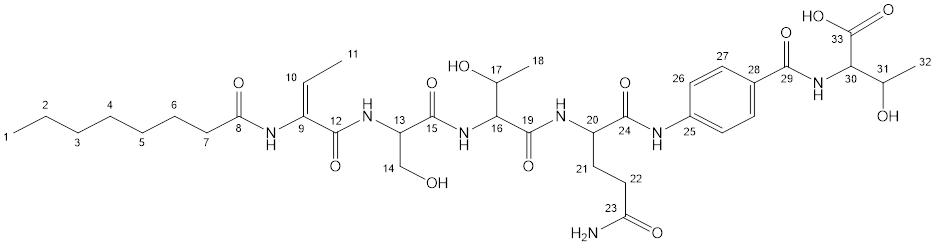

**A)** Key COSY, HMBC, and ROESY correlations in DMSO-d6 used to elucidate the structure of haereoacidicolin. **B)** Planar structure for haereoacidicolin with C-atom numbering.

### **Figure S7. ^1^H NMR spectrum for haereoacidicolin in DMSO-d_6_ at 600 MHz.**

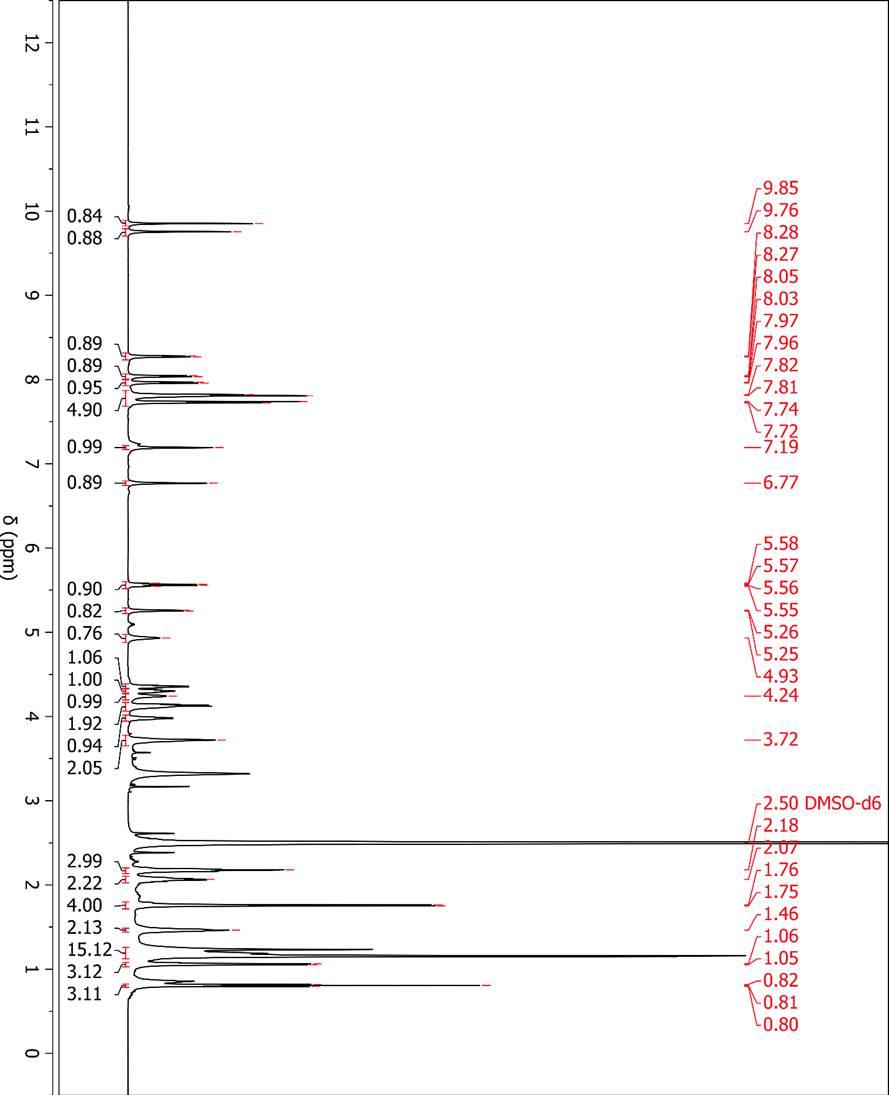

### **Figure S8. ^13^C NMR spectrum for haereoacidicolin in DMSO-d_6_ at 150 MHz.**

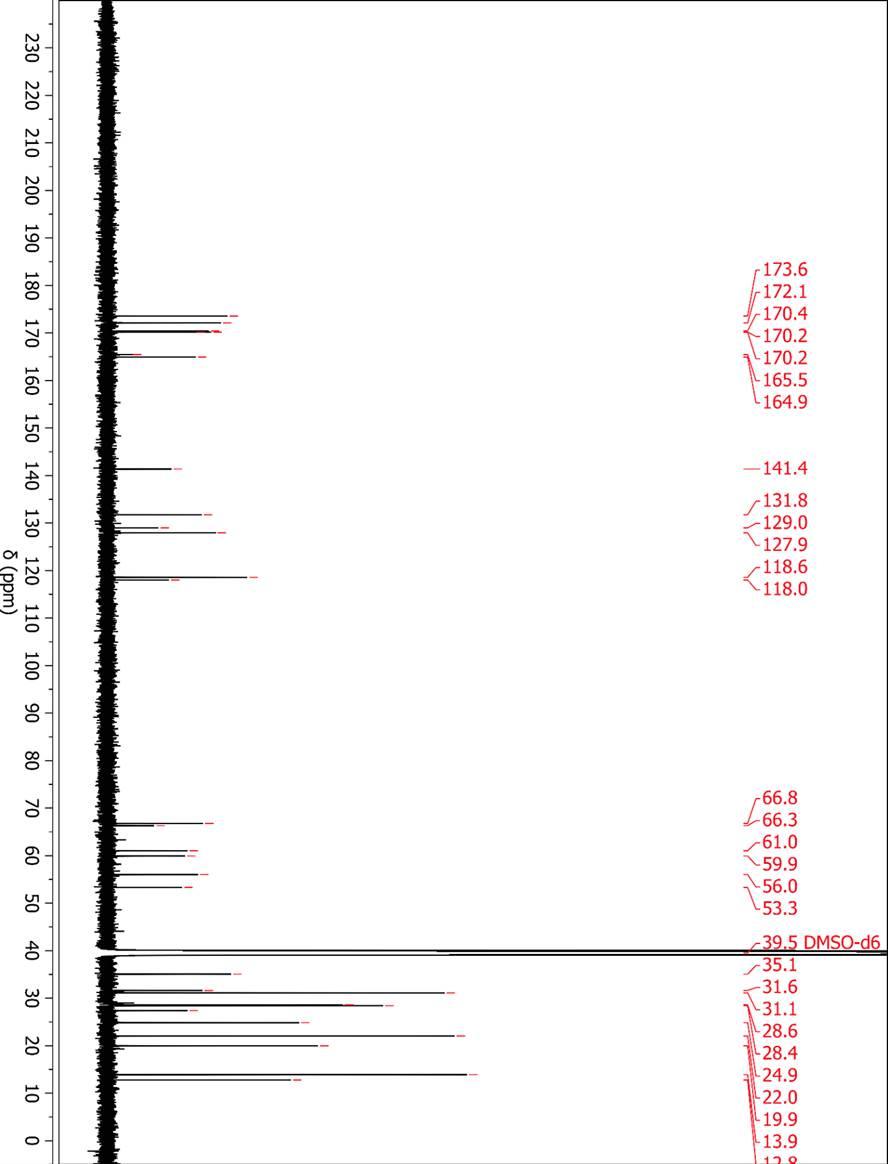

### **Figure S9. ^1^H-^13^C HSQC NMR spectrum for haereoacidicolin in DMSO-d_6_ at 600 MHz.**

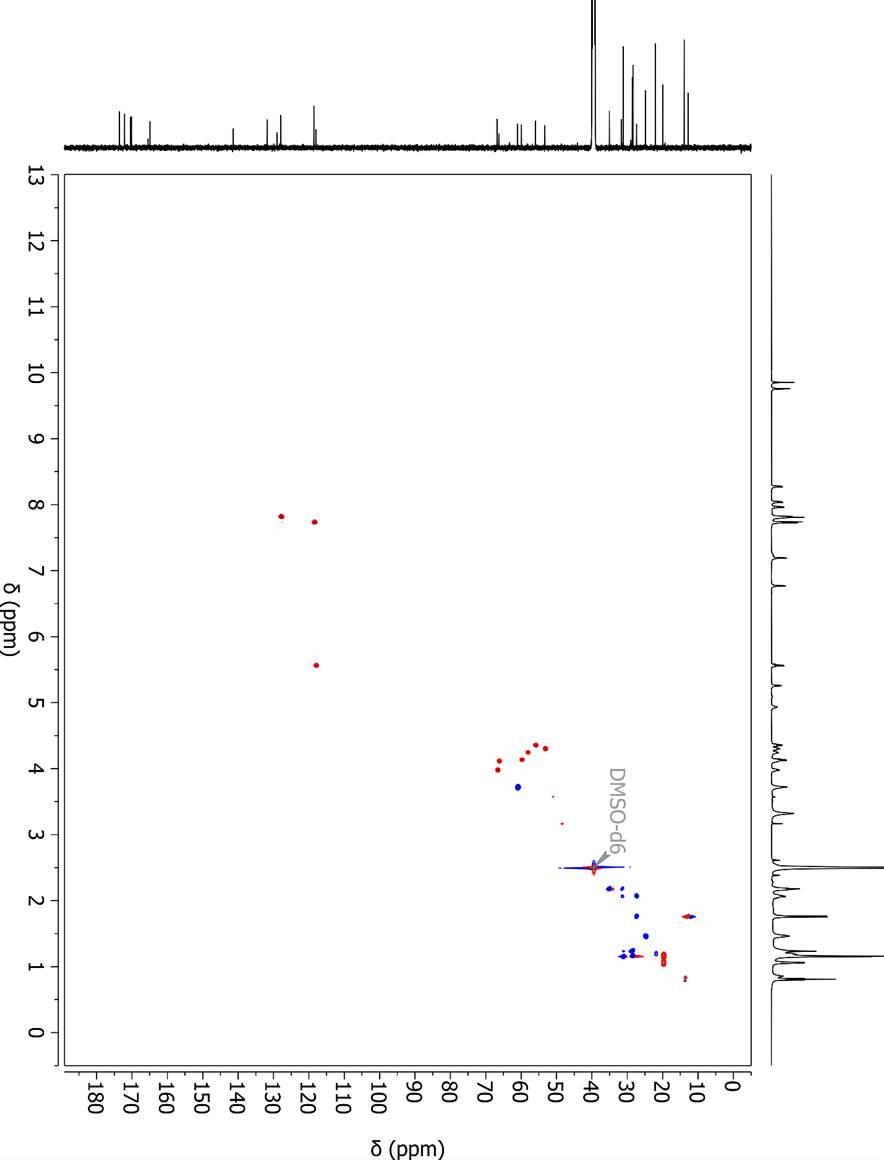

### **Figure S10. ^1^H-^1^H COSY NMR spectrum for haereoacidicolin in DMSO-d_6_ at 600 MHz.**

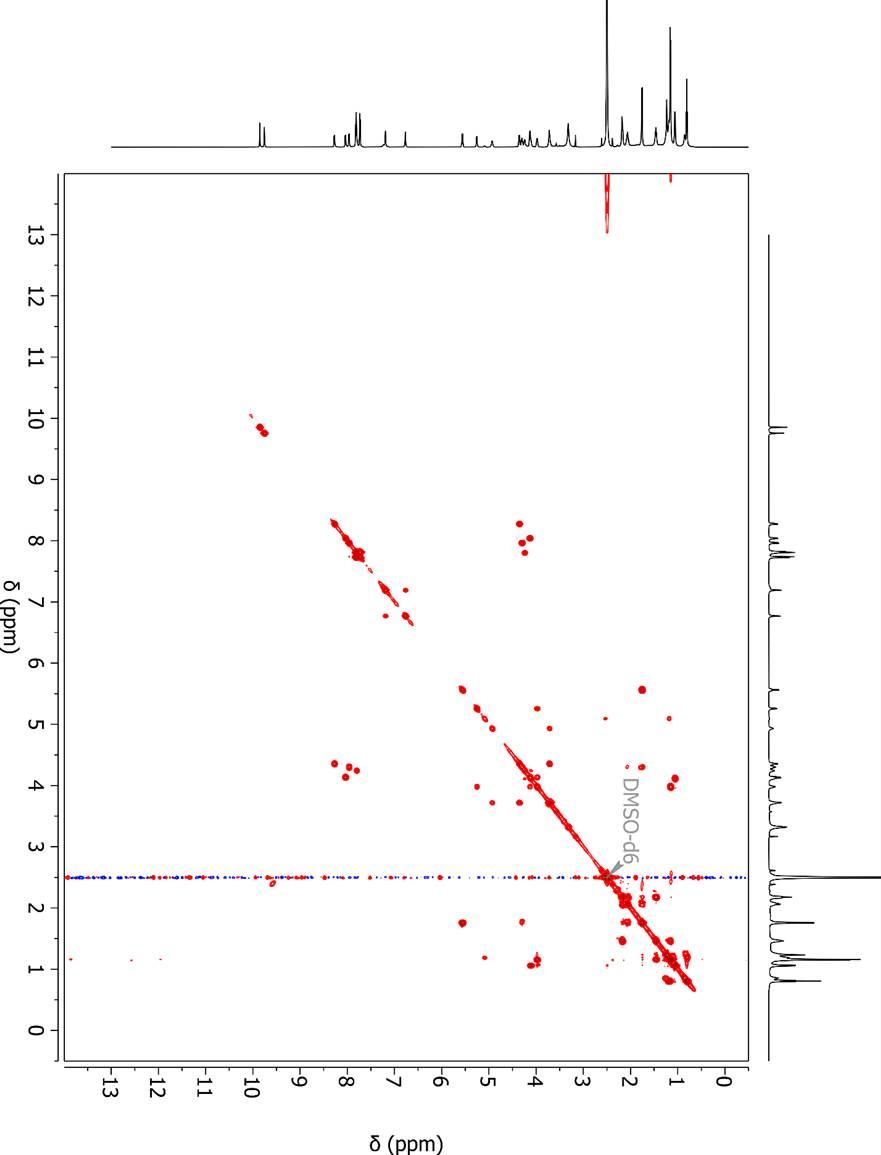

### **Figure S11. ^1^H-^13^C HMBC NMR spectrum for haereoacidicolin in DMSO-d_6_ at 600 MHz.**

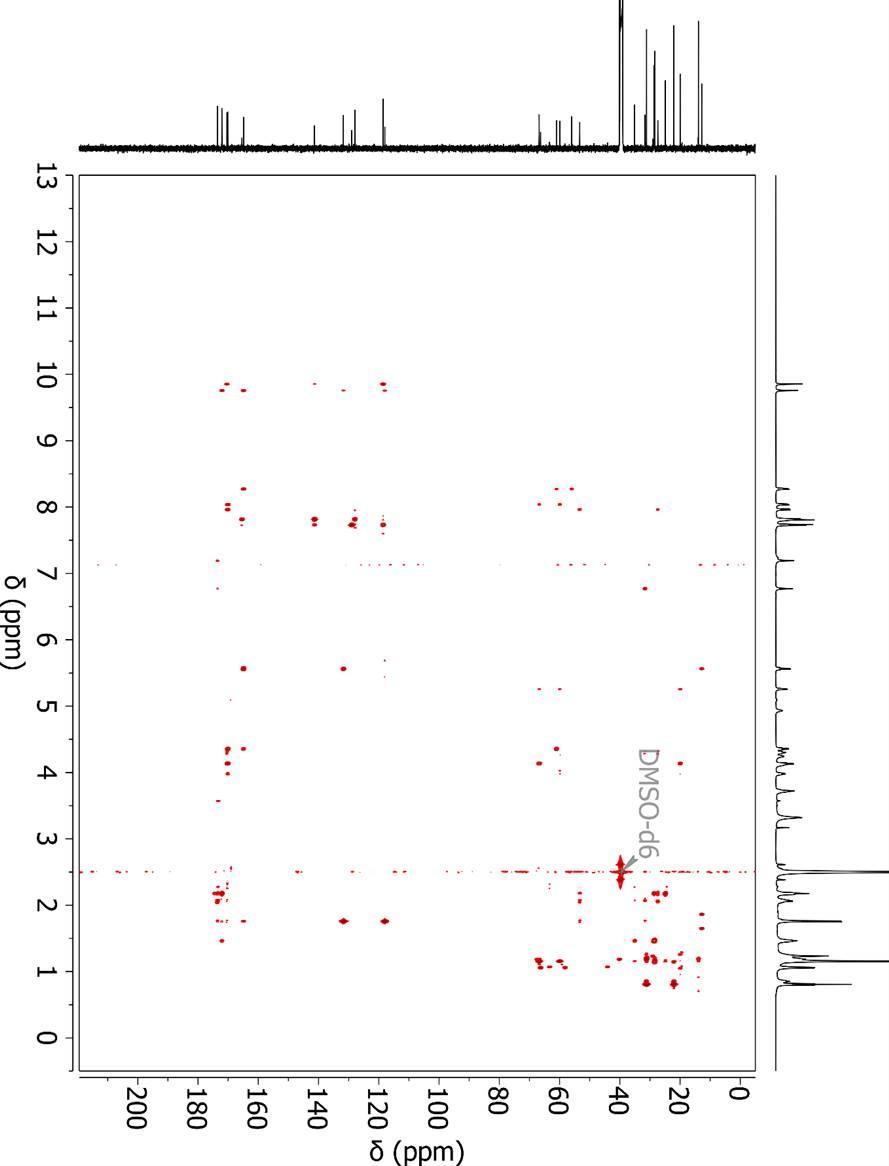

### **Figure S12. ^1^H-^1^H TOCSY NMR spectrum for haereoacidicolin in DMSO-d_6_ at 600 MHz.**

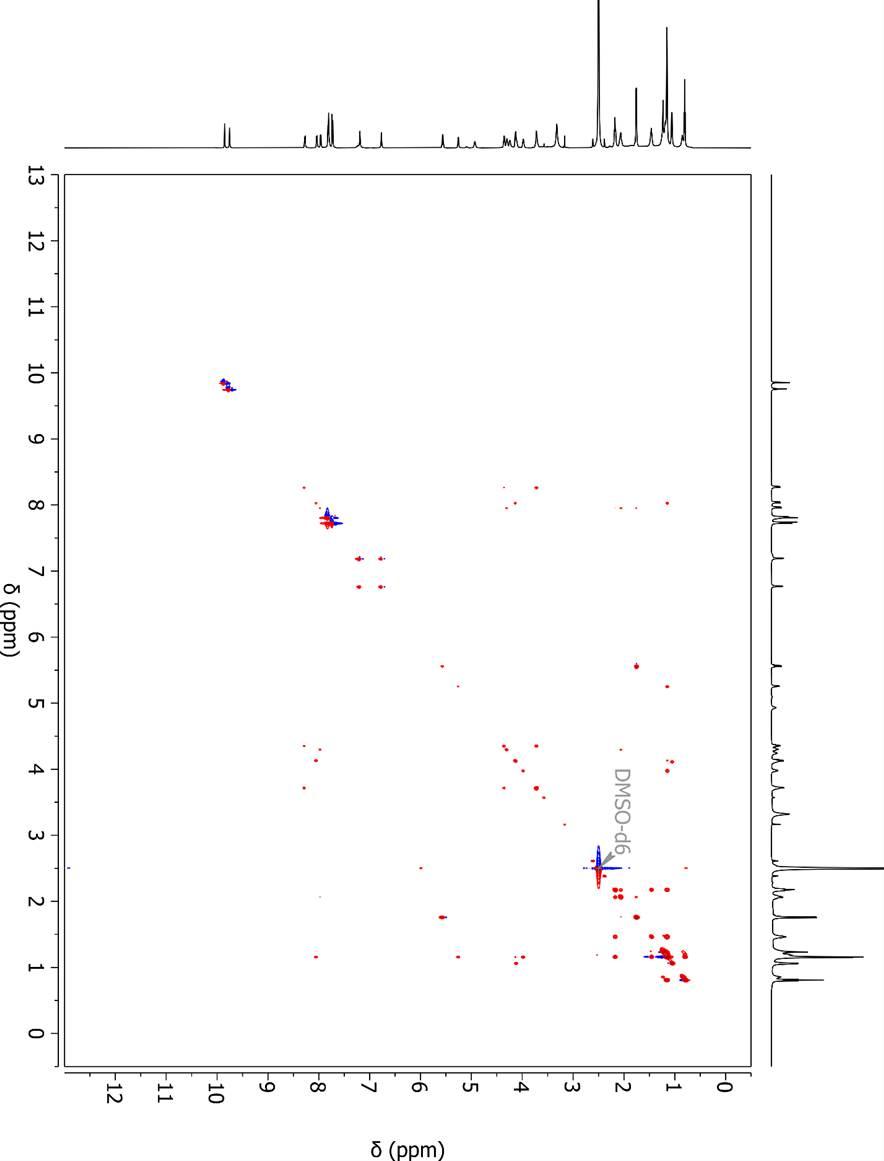

### **Figure S13. ^1^H-^15^N HSQC NMR spectrum for haereoacidicolin in DMSO-d_6_ at 600 MHz.**

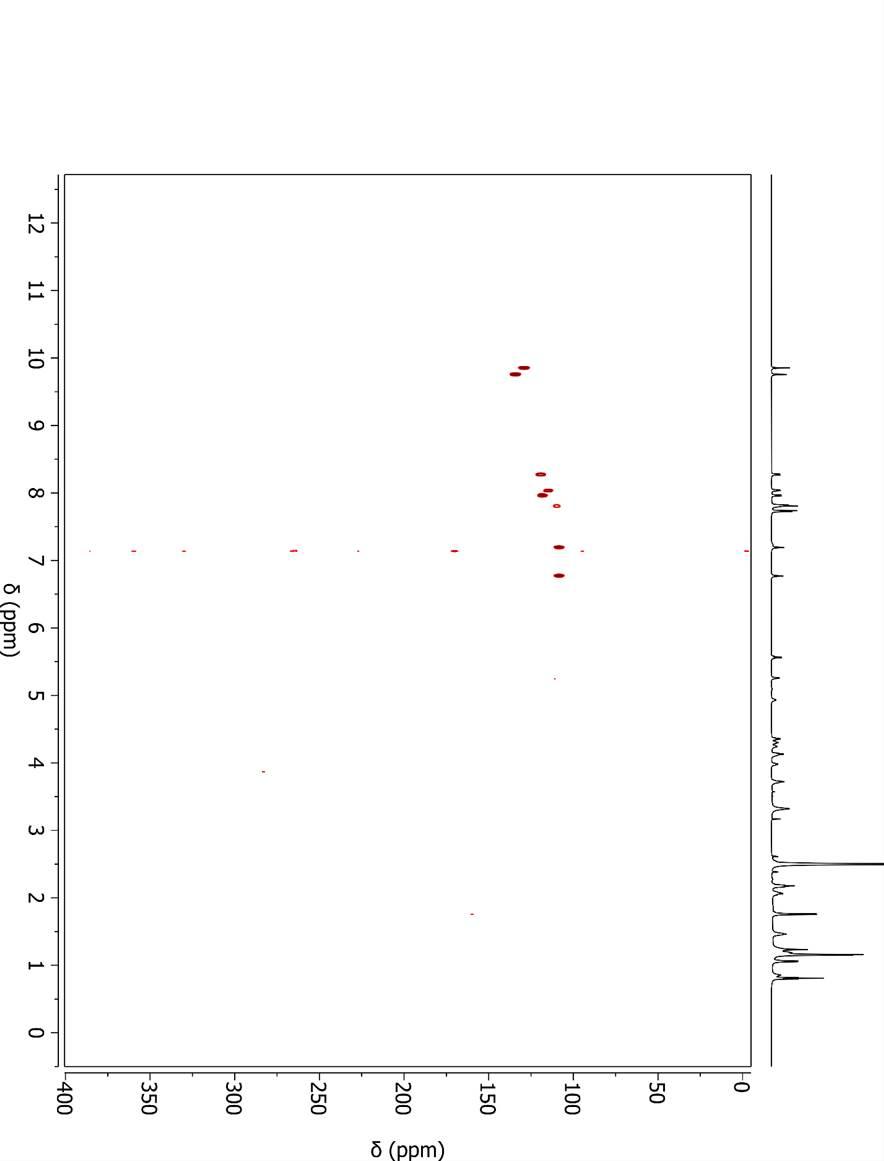

### **Figure S14. 1D selective ROESY NMR spectrum, irradiating 12-NH (δ_H_ 8.28), of haereoacidicolin in DMSO-d_6_ at 600 MHz.**

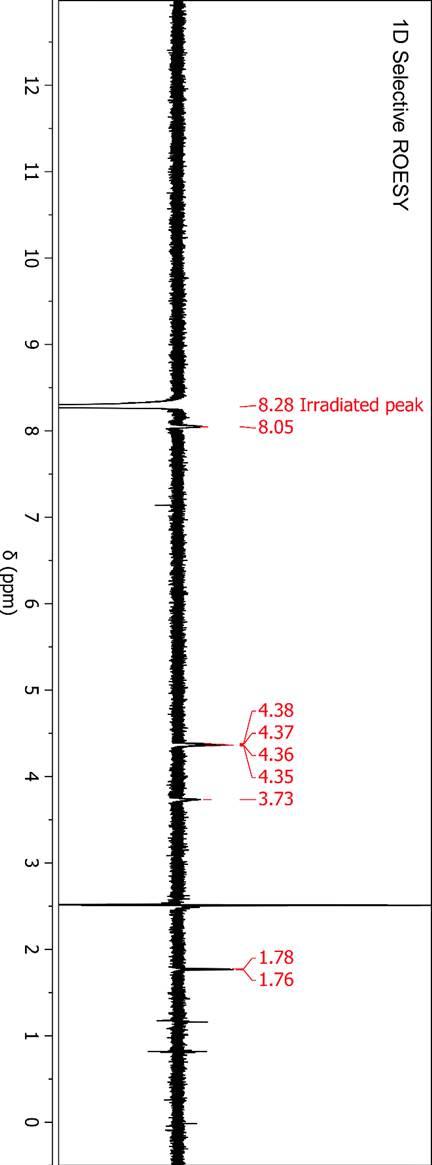

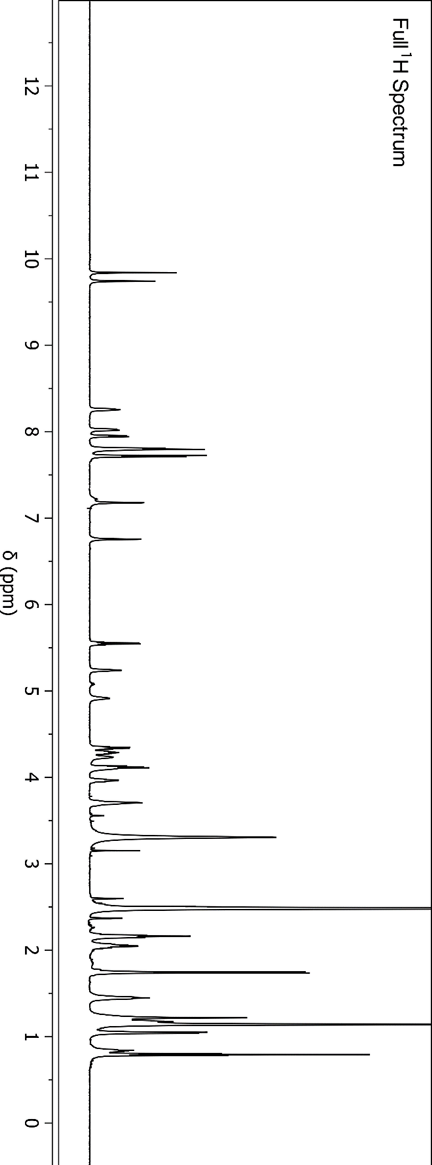

### **Figure S15. 1D selective ROESY NMR spectrum, irradiating 8-NH (δ_H_ 9.76), of haereoacidicolin in DMSO-d_6_ at 600 MHz.**

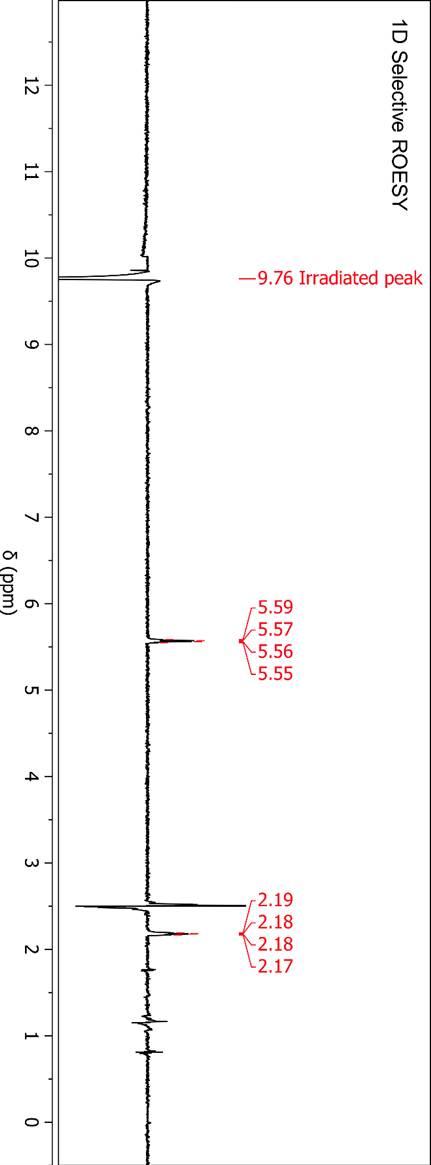

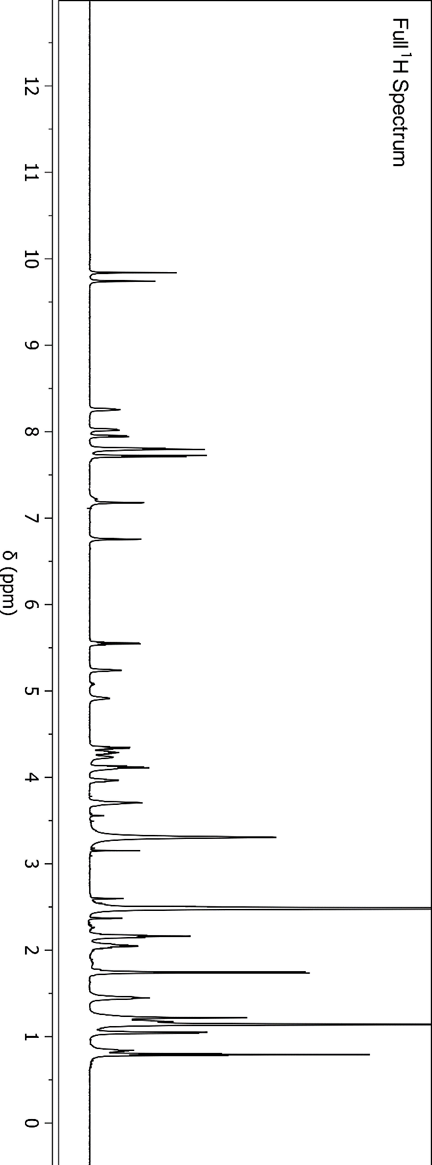

### **Figure S16. Extracted ion chromatogram for Marfey’s FDVA-glutamic acid hydrolysate.**

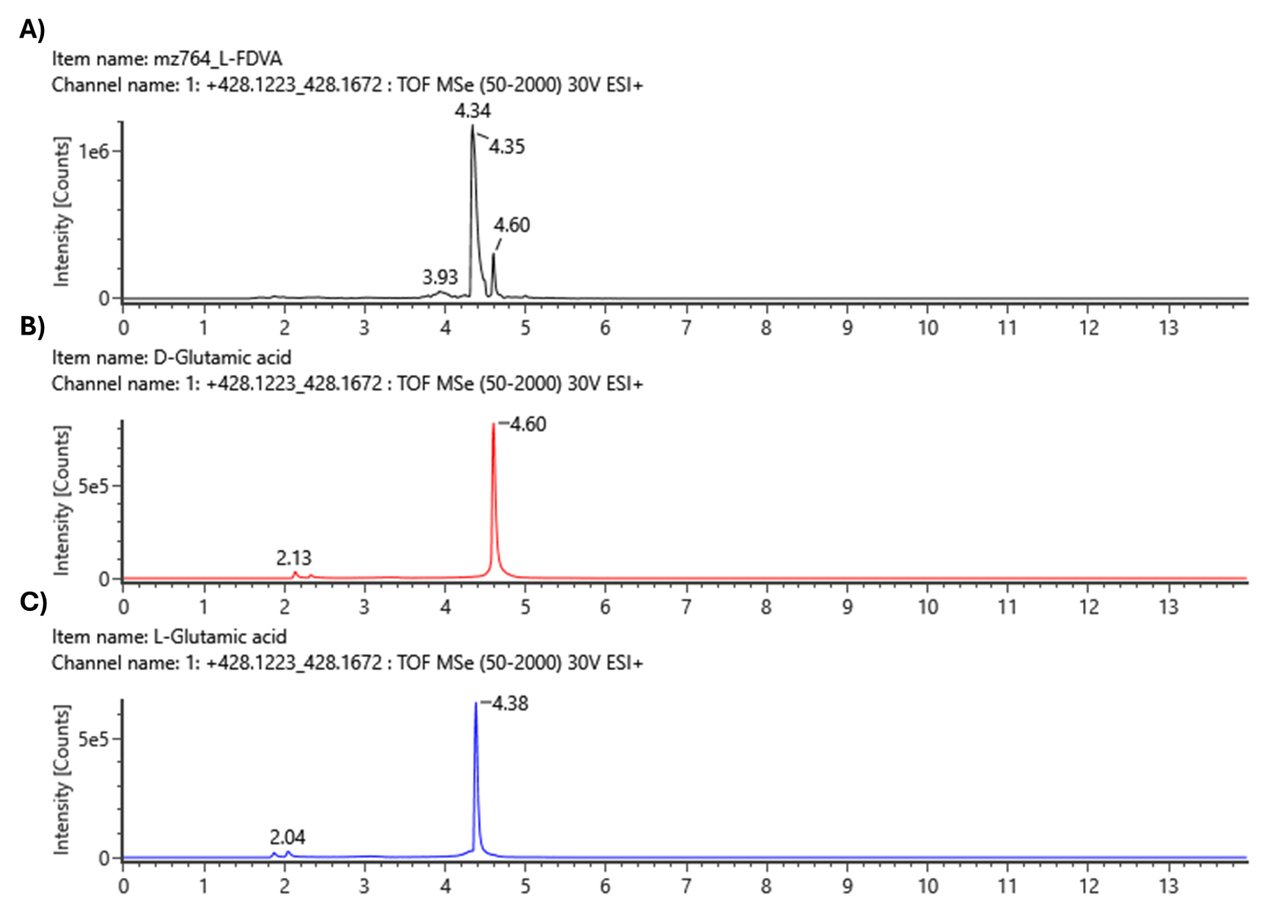

**(A)** Haereoacidicolin derivatized hydrolysate EIC *m/z* 428.1223–428.1672. **(B)** FDVA-ᴅ-glutamic acid EIC *m/z* 428.1223–428.1672; RT = 4.60 min. **(C)** FDVA-ʟ-glutamic acid EIC *m/z* 428.1223–428.1672; RT = 4.38 min.

### **Figure S17. Extracted ion chromatogram for Marfey’s FDVA-serine hydrolysate.**

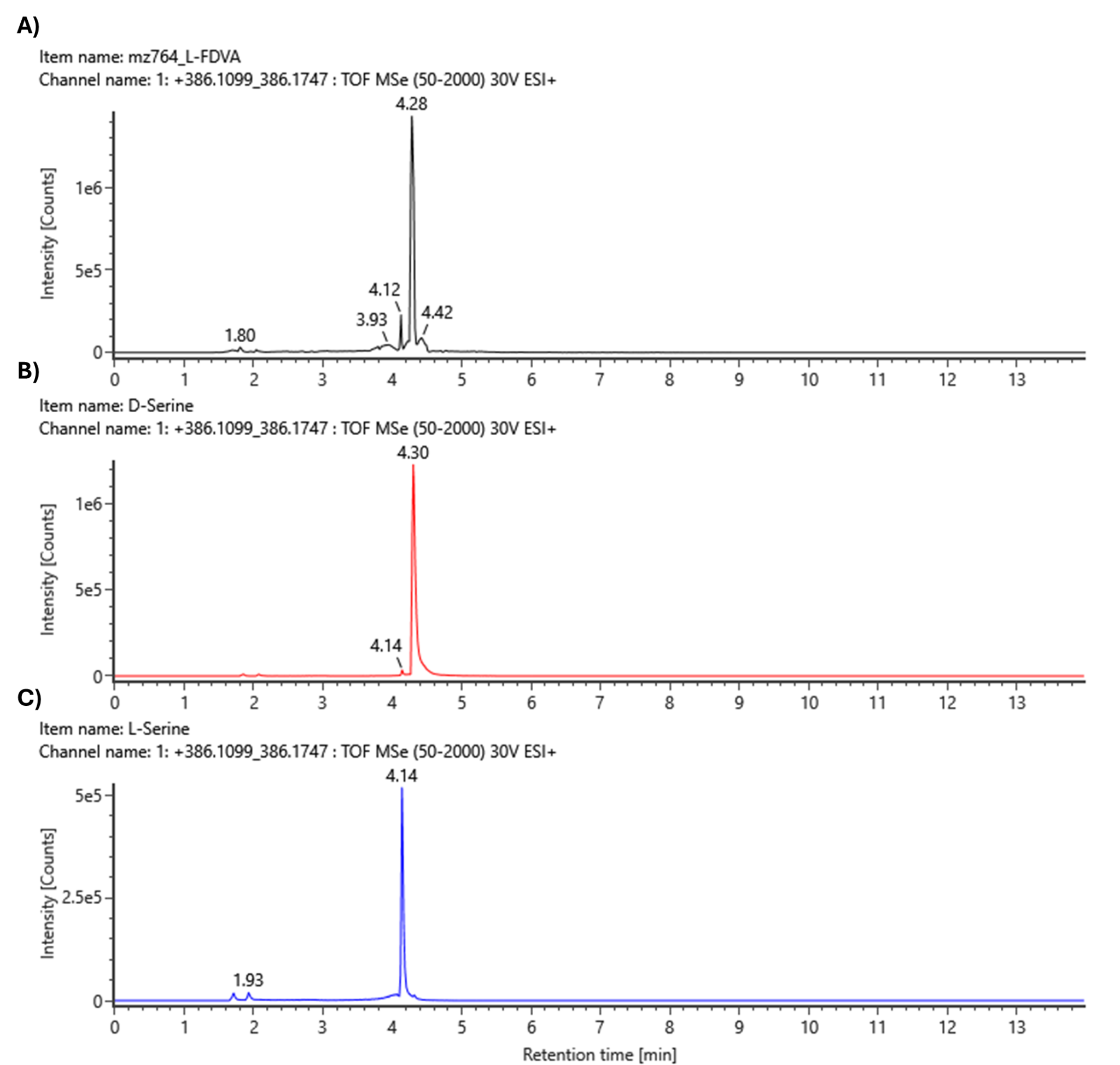

**(A)** Haereoacidicolin derivatized hydrolysate EIC *m/z* 386.1099–386.1747. **(B)** FDVA-ᴅ-serine EIC *m/z* 386.1099–386.1747; RT = 4.30 min. **(C)** FDVA-ʟ-serine EIC *m/z* 386.1099–386.1747; RT = 4.14 min.

### **Figure S18. Extracted ion chromatogram for Marfey’s FDVA-threonine hydrolysate.**

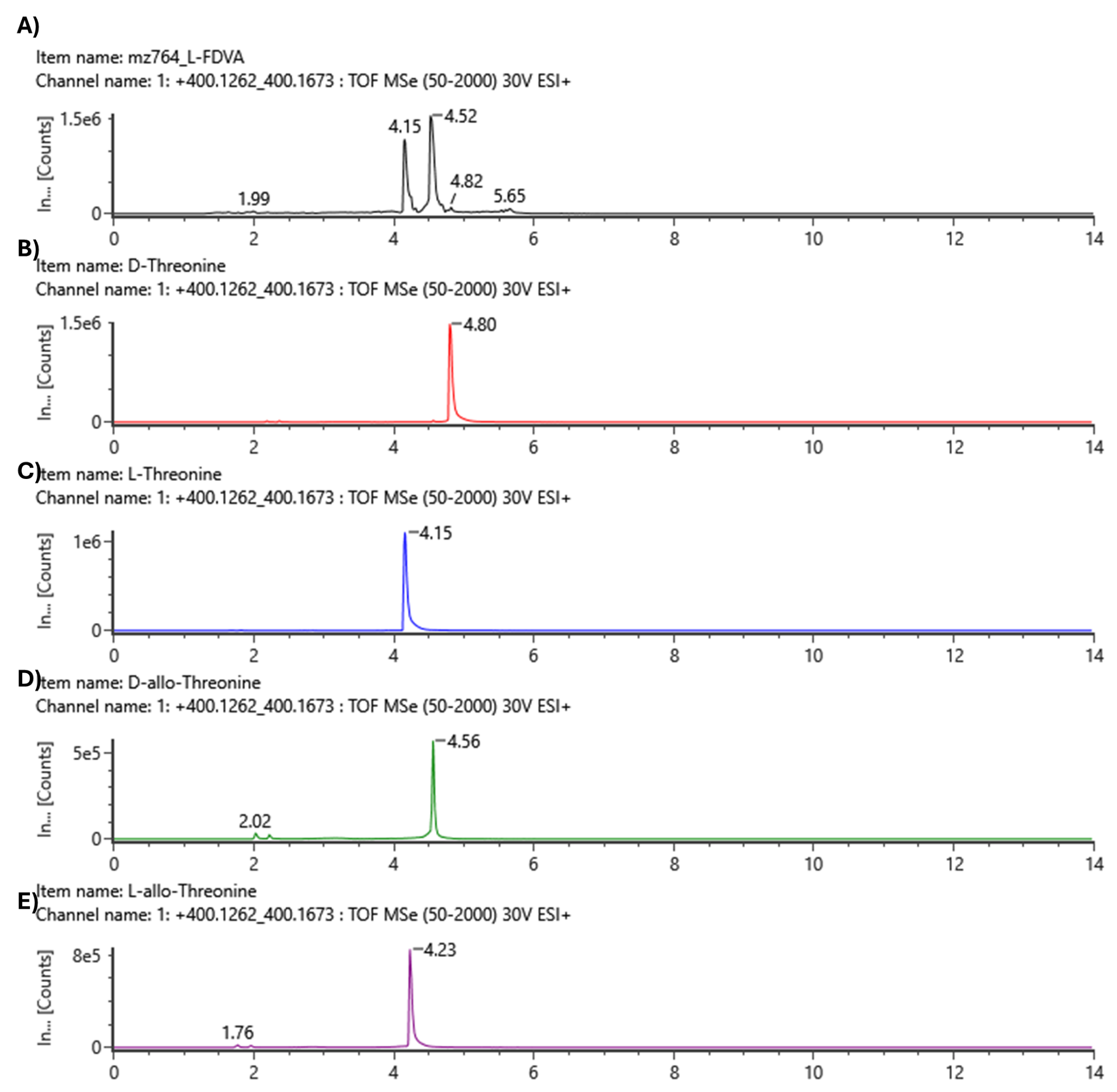

**(A)** Haereoacidicolin derivatized hydrolysate EIC *m/z* 400.1262–400.1673. **(B)** FDVA-ᴅ- threonine EIC *m/z* 400.1262–400.1673; RT = 4.80 min. **(C)** FDVA-ʟ-threonine EIC *m/z* 400.1262–400.1673; RT = 4.15 min. **(D)** FDVA-ᴅ-allo-threonine EIC *m/z* 400.1262–400.1673; RT = 4.56 min. **(E)** FDVA-ʟ-allo-threonine EIC *m/z* 400.1262–400.1673; RT = 4.23 min.

### **Figure S19. Neighbor-joining phylogenetic tree of 189 validated NRPS condensation domains.**

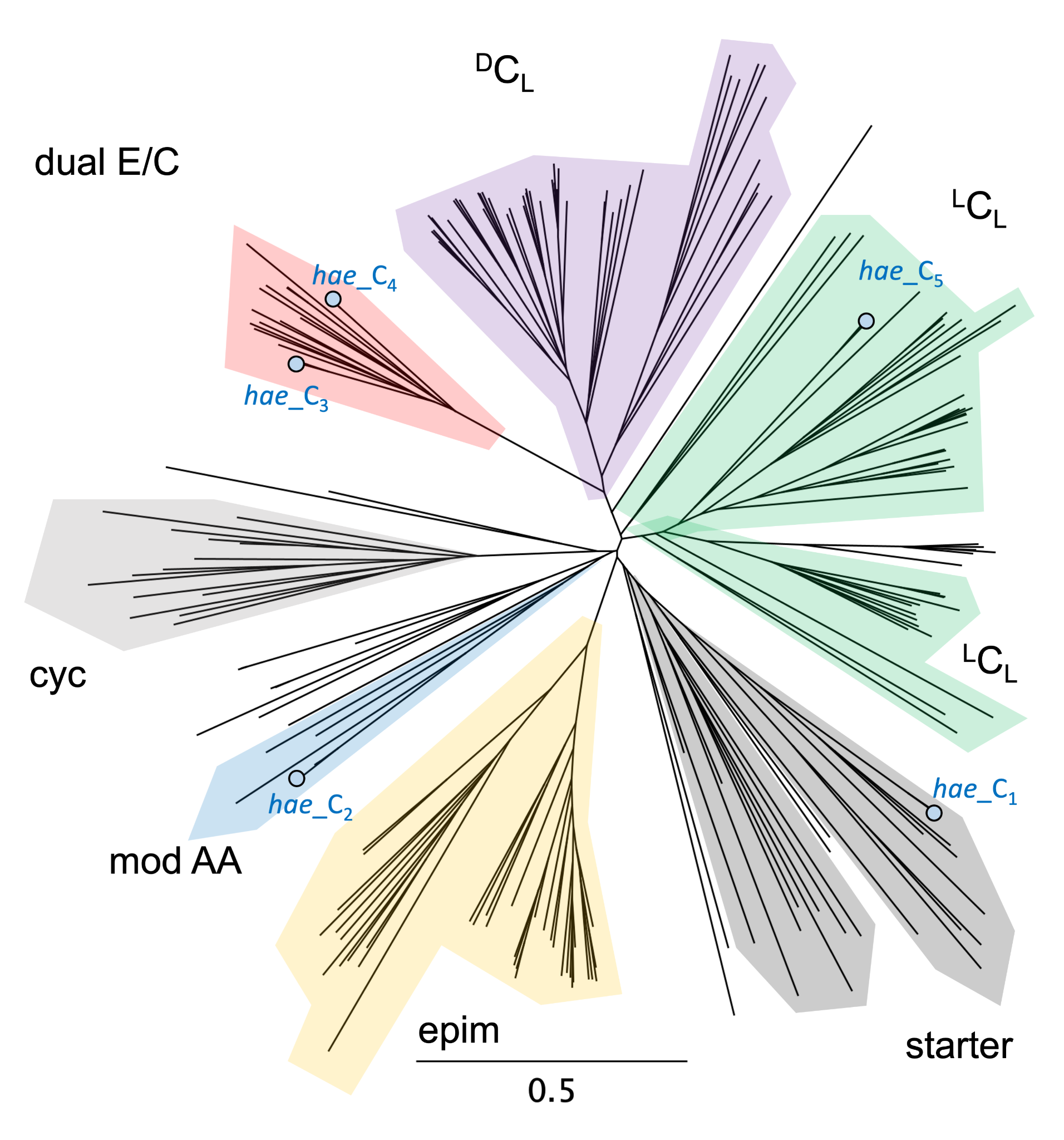

Blue dots represent the 5 *hae* condensation (C) domains. The C domains from *P. megapolitana* 3-E9, *P. acidicola* 1-D8, and *P. megapolitana* DSM 23488 clade closely together and therefore are represented by a single dot. The phylogeny of C domains are consistent with proposed functional classification and offer a comparative framework to support experimental data. The phylogenetic tree was generated using Geneious Prime (2025.1.3) software with the JTT model of amino acid substitution. The scale bar represents the average number of amino acid substitutions. Starter, starter C domains are associated with acylation of the first amino acid. ^L^C_L_, C domains that catalyze formation of a peptide bond between two ʟ-amino acids. ^D^C_L_, C domains that condense a d-amino acid donor with an l-amino acid acceptor. Cyc, cyclization domains that catalyze both peptide bond formation and subsequent cyclization of cysteine, serine or threonine residues. Epim, epimerization domain. Dual E/C, C domains that catalyze epimerization and condensation. Mod AA, C domains that catalyze peptide bond formation and modification of the incorporated amino acid such as dehydration of threonine to Dhb.

### **Supplementary Note 4. Multi-Lab Utility.**

In order to demonstrate cross user/instrument utility, we selected all 26 *B. subtilis* spectra from the knowledgebase, which span three laboratories and five cultivation and analysis protocols. Using the introduced intensity-agnostic cosine method, pairwise spectral distances showed statistically significant differences in fingerprints between *B. subtilis* and other *Bacillus* spp. (n=24 spectra) (PERMANOVA, pseudo-F = 7.51, p = 0.001). Notably, PERMDISP analysis failed to support the conclusion that inter-group variance was significantly different (F = 2.60, p = 0.08) suggesting that the observed separations likely reflect differences in centroid locations. Further, inter- and intra- species distances were quantified revealing that the average distance within *B. subtilis* and other *Bacillus* spp. increases from 0.6821 to 0.8316 (**Fig. S20A**). Most notably, for all *B. subtilis* strains, their nearest neighbors were intra-species. To visually confirm the result, PCoA analysis was performed (reference **Fig. S20 B**) revealing that *B. subtilis* strains formed two distinct clusters with four outliers (further analysis below). Together, these findings highlight that the IDBac analysis platform performed well for all three labs, however inter-laboratory variation was observed.

To further explore the variation observed within *B. subtilis* spectra shown in **Fig. S20** in addition to factors driving inter-lab reproducibility, we again performed PCoA analysis and visualized the results in the context of the metadata already available in the IDBac Knowledgebase (**Fig. S21**). As expected (DOI: 10.1080/19490976.2020.1740073), cultivation and MALDI preparation protocols are paramount in replicability. Even within the same lab, across strains differences in proteomic profiles could be observed. Nevertheless, in a database search scenario, the nearest neighbor for all strains was found to be within-species, highlighting the efficacy of the platform for multiple laboratories. Through the continued growth of the IDBac knowledgebase as a resource, an increasing number of protocols will be covered thus enabling broad utility across laboratories and preparation procedures. In the future, the collection of such data will enable a large-scale analysis of protocols and their effects on microbial identification, further advancing biotyping for the broader community.

Additionally, as mentioned in the main text, 16S rRNA gene analysis can be limited to genus level assignments. For all thirteen strains that were identified through 16S rRNA gene sequencing analysis, a nucleotide BLAST search was performed, retrieving the top 100 candidates. Results were filtered for an e-value of 0.0, an identity >99%, and a species-level annotation. In all cases the percent of results that were *B. subtilis* did not exceed 75%, and as a result, the possibility that these spectra reflect a different species cannot be ruled out. The platform continuously checks for updates in classification from the genomic identifier and will be updated should a change occur. Variability in proteomic fingerprints controlling for this factor are further evaluated in **Supplementary Note 5**.

### **Figure S20. *Bacillus* spp. similarities.**

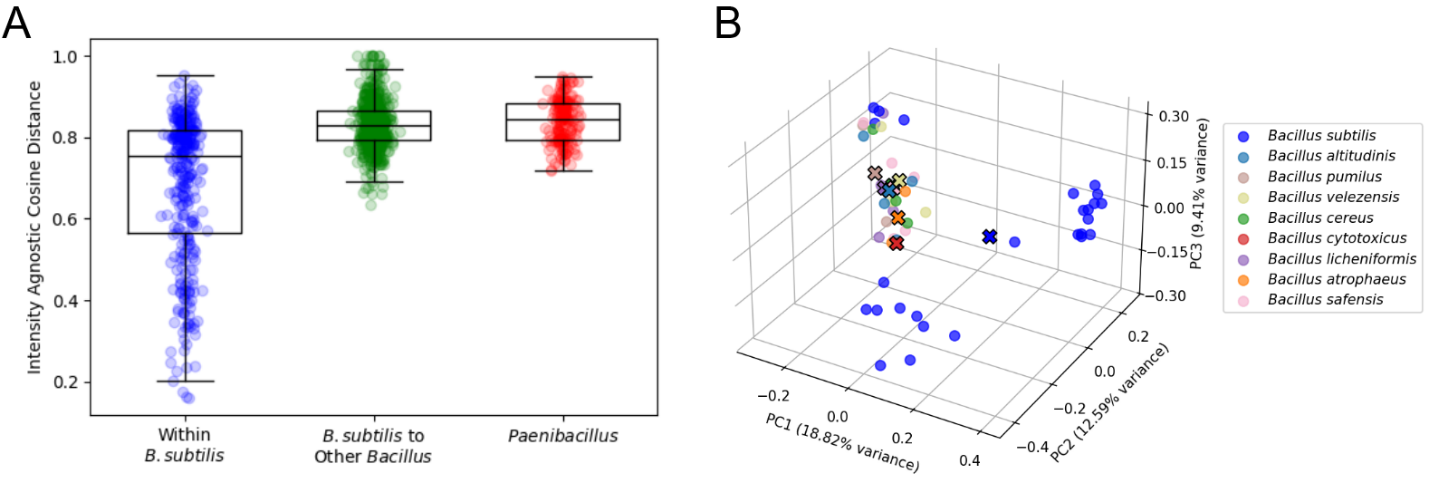

***Bacillus* spp. similarities.** **A.** Distribution of pairwise similarities for within *B. subtilis* (left), between *B. subtilis* and other *Bacillus* spp. (middle), and strains from the closely related *Paenibacillus* genus (right). A statistically significant difference in the multivariate profiles between *B. subtilis* and other *Bacillus* taxa was observed (PERMANOVA, p = 0.001). PERMDISP did not support a significant difference in dispersion between groups (p = 0.08) suggesting that centroid location was the primary factor in the PERMANOVA result. **B**. PCoA projection of all *Bacillus* strains with centroids denoted by ‘X’ markers. In all cases, nearest neighbors of *B. subtilis* strains are other *B. subtilis* spectra in cosine space.

### **Figure S21. Cross-laboratory analyses.**

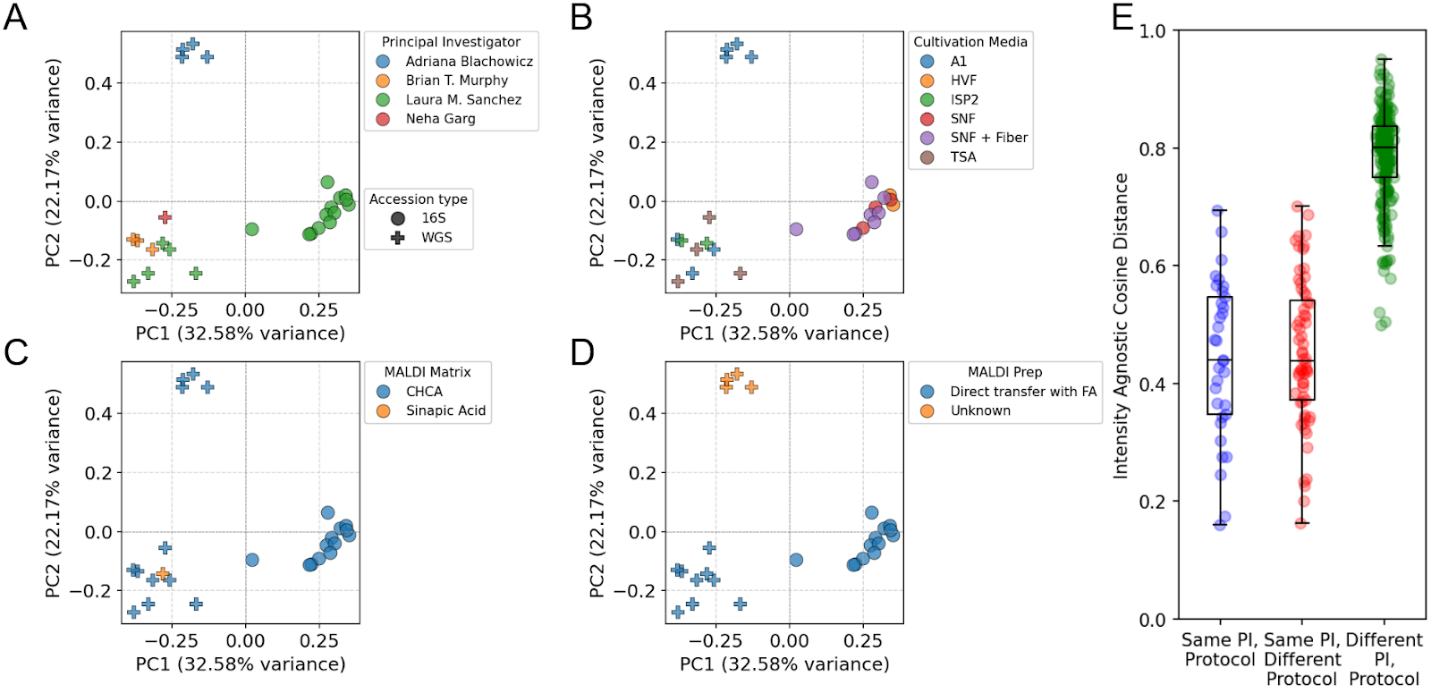

*PCoA analysis of B. subtilis strains in the IDBac Knowledgebase across laboratories*. Genomic confirmation for each strain is highlighted by shape. Data points are colored by **A)** Principal investigator, **B)** Cultivation media **C)** MALDI matrix, and **D)** Cell mass transfer technique. Differences between sample cultivation and preparation procedures have observable differences in proteomic profiles. **E)** When controlling for PI, cultivation media, matrix, and transfer technique, MALDI-TOF data sets are reproducible.

### **Supplementary Note 5.** **Strain Reproducibility.**

To evaluate reproducibility in a controlled setting, we measured the spectral similarity of *B. subtilis* 3610 strains across laboratories and instruments, controlling for cultivation conditions. Specifically, two *B. subtilis* 3610 cryovials (created and dispersed to each lab nine years prior) were retrieved from -80 °C storage and cultured on A1 and TSA media for 48 hours at 25 °C. Spectra were collected by two independent users on Bruker Daltonics microflex and rapifleX instruments resulting in a standard cosine distance score of 0.23 and 0.36 respectively (intensity agnostic cosine = 0.55, 0.60) demonstrating the reproducibility of the approach (**Fig. S22A-B**).

To demonstrate the consistency of our approach for a given strain, independent of cultivation and MALDI preparation protocols, an aggregated analysis was performed using all nine *B. subtilis* 3610 strains currently available in the KB. In a cross-laboratory query with the intensity agnostic distance metric, 7/9 top-1 hits were annotated as *B. subtilis* 3610 strains, and 100% of the top-1 and top-3 hits for all nine strains were within *B. Subtilis*.

An evaluation of distances from *B. subtilis* 3610 to other KB strains using both the intensity agnostic cosine distance (**Fig. S22C**) and standard cosine distance (**Fig. S22D**) empirically shows that within-strain distances are lower than species- and genus-level hits, highlighting the reproducibility of this method across laboratories and protocols. Further, this analysis suggests that the intensity agnostic cosine distance may reduce false-positives in genus-level identifications relative to the standard cosine score, however a larger scale analysis is needed to confirm this.

### **Figure S22.** **Reproducibility**

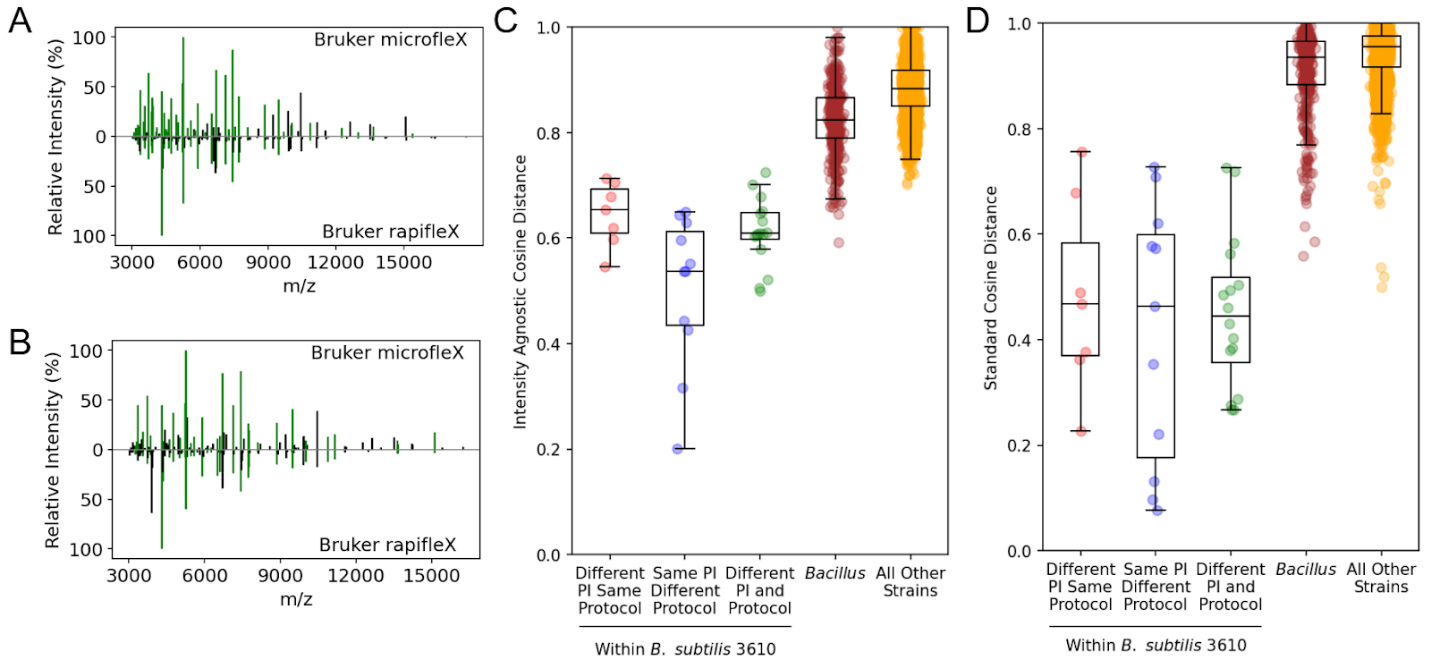

**A** & **B** Mirror plots demonstrating the reproducibility of *B. subtilis* 3610 strains cultivated on A1 (**A**) and TSA (**B**) media across laboratories and instruments. **C&D** Analysis of distances between *B. subtilis* 3610 strains controlling for protocol and laboratory. Distance to other strains within the same genus and all other KB strains is also shown highlighting the distinction in distances between strain-level hits and genus-level hits. **C.** The intensity agnostic cosine better differentiation at the genus level relative to standard cosine (**D**).

### **Table S5. Initial IDBac-KB contributions by lab/institution.**

| **Laboratory / Institution** | **# of spectra contributed** | **# of unique genera** |
| --- | --- | --- |
| Adaikpoh Laboratory  University of Illinois Chicago | 8 | 5 |
| Bugni Laboratory  University of Wisconsin Madison | 140 | 4 |
| Pamer Laboratory  Duchossois Family Institute University of Chicago | 156 | 16 |
| Federle Laboratory  University of Illinois Chicago | 19 | 2 |
| Garg Laboratory  Georgia Institute of Technology | 14 | 7 |
| Henke Laboratory  University of Illinois Chicago | 18 | 8 |
| Jensen Laboratory  Scripps Institution of Oceanography  University of California San Diego | 11 | 2 |
| Linington Laboratory  Simon Fraser University | 92 | 9 |
| Murphy Laboratory  University of Illinois Chicago | 214 | 68 |
| NASA Jet Propulsion Laboratory | 57 | 12 |
| NASA Marshall Space Flight Center | 54 | 24 |
| Stevens Laboratory  University of Mississippi | 1 | 1 |
| USDA-ARS NRRL & Metcalf Laboratory  University of Illinois Urbana-Champaign | 419 | 9 |
| Sanchez Laboratory  University of California Santa Cruz | 62 | 19 |
| Winter Laboratory  University of Utah | 10 | 1 |
| McCauley and Carlson Laboratories  California State University Dominguez Hills  University of the Pacific | 28 | 7 |
| McKinnie Laboratory  University of California Santa Cruz | 3 | 2 |

### **Table S6. Decomposition study environmental sample conditions.**

| ^*^**Swab** | **Hours postmortem** | **Swab surface** | **Swab location temperature (°C)** | **Swab location pH** | **Ambient temperature (°C)** | **Relative humidity (%)** | **Sun exposure** |
| --- | --- | --- | --- | --- | --- | --- | --- |
| 1 | 6 | Skin; right shoulder | 32.5 | 4.3 | 27.1 | 60 | Direct sun all day |
| 2 | 30 | Skin; right shoulder | 36.9 | 4.3 | 28.8 | 51 | Direct sun all day |
| 3 | 78 | Skin; right shoulder | 34.0 | 5.7 | 27.4 | 54 | Direct sun all day |
| 4 | 102 | Skin; right shoulder | 37.4 | 5.9 | 28.0 | 52 | Direct sun all day |

*Samples used in this analysis were obtained from the laboratory of Dr. David Carter, while at the Chaminade University of Honolulu. All carcasses were acquired commercially (Island Farms, Waianae, USA) and handled in accordance with State of Hawaii Statute §159–21.

### **Table S7. Decomposition study media ingredients.**

| **Media type** | **Ingredient** | **Weight/vol (g/L)** |
| --- | --- | --- |
| **A1** | Soluble starch | 10 g |
|  | Yeast extract | 4 g |
|  | Peptone | 2 g |
|  | Agar | 15 g |
|  | Sterilized dH_2_O | 1000 mL |
| **M1** | Soluble starch | 10 g |
|  | Yeast extract | 4 g |
|  | Peptone | 2 g |
|  | Agar | 15 g |
|  | Sterilized dH_2_O | 1000 mL |
| **R2A** | Yeast extract | 0.5 g |
|  | Meat peptone | 0.5 g |
|  | Casamino acids | 0.5 g |
|  | Glucose | 0.5 g |
|  | Starch | 0.5 g |
|  | Dipotassium hydrogen phosphate | 0.3 g |
|  | Magnesium sulphate | 0.05 g |
|  | Sodium pyruvate | 0.3 g |
|  | Agar | 15 g |
|  | Sterilized dH_2_O | 1000 mL |
| **M1+** | Soluble starch | 5 g |
|  | Yeast extract | 2 g |
|  | Peptone | 1 g |
|  | Agar | 8 g |
|  | Gellan gum | 8 g |
|  | Sterilized dH_2_O | 1000 mL |

### **Table S8. Cave system isolation-media ingredients.**

| **Media type** | **Ingredient** | **Weight/vol (g/L)** |
| --- | --- | --- |
| **A1** | Soluble starch | 10 g |
|  | Yeast extract | 4 g |
|  | Peptone | 2 g |
|  | Agar | 15 g |
|  | Sterilized dH_2_O | 1000 mL |
| **M31**  Adapted from Ivanova et al., 2016^5^ | KH2PO4 | 0.1 g |
|  | Hutner’s basal salts:  Nitrilotriacetic acid (NTA), MgSO4. 7H20, CaCl2, 2H2O, (NH4) MoO7O24. 4H2O, FeSO4. 7H2O    “Metal 44”:  Na-EDTA, ZnSO4. 7H2O, FeSO4. 7H2O, MnSO4. H2O, CuSO4. 5H2O, Co(NO3)2. 6H2O, Na2B4O7. 10H2O, Distilled water | 20 mL |
|  | N-acetylglucosamine | 1.0 g |
|  | Peptone | 0.1 g |
|  | Yeast extract | 0.1 g |
|  | Gellan gum | 8.00 g |
|  | Agar | 8.00 |
|  | dH_2_O | 980 mL |
| **629**  As described by Leibniz Institute DSMZ-German Collection of Microorganisms and Cell Cultures. | Peptone | 5.0 g |
|  | Yeast extract | 0.5 g |
|  | Hutner’s basal salts:  Nitrilotriacetic acid (NTA), MgSO4. 7H20, CaCl2. 2H2O, (NH4) MoO7O24. 4H2O, FeSO4. 7H2O  “Metal 44”:  Na-EDTA, ZnSO4. 7H2O, FeSO4. 7H2O, MnSO4. H2O, CuSO4. 5H2O, Co (NO3)2. 6H2O, Na2B4O7. 10H2O, Distilled water | 20.0 mL |
|  | Staley's vitamins:  Vitamin B12, Biotin, Thiamine-HCl x 2 H2O, Ca-pantothenate, Folic acid, Riboflavin, Nicotinamide & Distilled water | 10.0 mL |
|  | Gellan gum | 8.00 g |
|  | Agar | 8.00 |
|  | Distilled water | 970.0 mL |

*Samples used in this analysis were obtained from the laboratory of Dr. Hazel Barton, while at the University of Akron.

### **Table S9. MALDI-TOF MS instrument specifications and parameters for initial IDBac-KB populations.**

| **Instrument** | **Laser** | **Shots** | **RepRate** | **Delay** | **Ion source 1 voltage** | **Ion source 2 voltage** | **Lens voltage** | **Mass range** | **Matrix suppression cutoff** | **Software** |
| --- | --- | --- | --- | --- | --- | --- | --- | --- | --- | --- |
| Autoflex Speed LRF mass spectrometer (Bruker Daltonics) | smartbeam™-II (355 nm) | 1200 | 2000 Hz | 29731 ns | 19.5 kV | 18.35 kV | 7 kV | 1.92 kDa-  21 kDa | 1.9 kDa | flexAnalysis software  v. 3.4. |
| rapifleX MALDI Tissuetyper mass spectrometer (Bruker Daltonics) | smartbeam™ 3D laser  (355 nm) | 2000 | 5000 Hz | 28272 ns | 20 kV | 18.45 kV | 9 kV | 1.80 kDa-  21 kDa | 1.6 kDa | flexAnalysis software  v. 4.0.46.0 |
| microflex LT MALDI-TOF mass spectrometer (Bruker Daltonics) | class III B nitrogen laser (337 nm) | 600 | 60 Hz | 150 ns | 20kV | 18.13 kV | 6.02kV | 2 kDa- 20 kDa | N/A | flexAnalysis software  v. 3.4  (Build 79) |
| 8020 Benchtop MALDI-TOF mass spectrometer  (Shimadzu) | ND:YAG  laser (355 nm) | 300 | 200 Hz | 102 ns | 20kV | 1.1 kV | - | 1.9kDa-21kDa | N/A | MALDI solutions data acquisition software  v 2.8.0 |

### **Figure S23.** **Quality control of protein MS spectra for IDBac-KB depositions.**

**Unprocessed MALDI MS data from three strains representing criteria that did not meet data cleanliness requirements. A)** Proteomic fingerprints fall into two distinct groupings (replicates 1-3, and 4-10) suggesting that more than one strain may be present among replicates. **B)** No consistent peaks at a signal to noise ratio greater than 4 occurred in greater than 50% of replicates, and **C)** Replicates one and two show noisy baselines may impact peak picking from the raw data. Replicates one and three show lower intensity than anticipated.

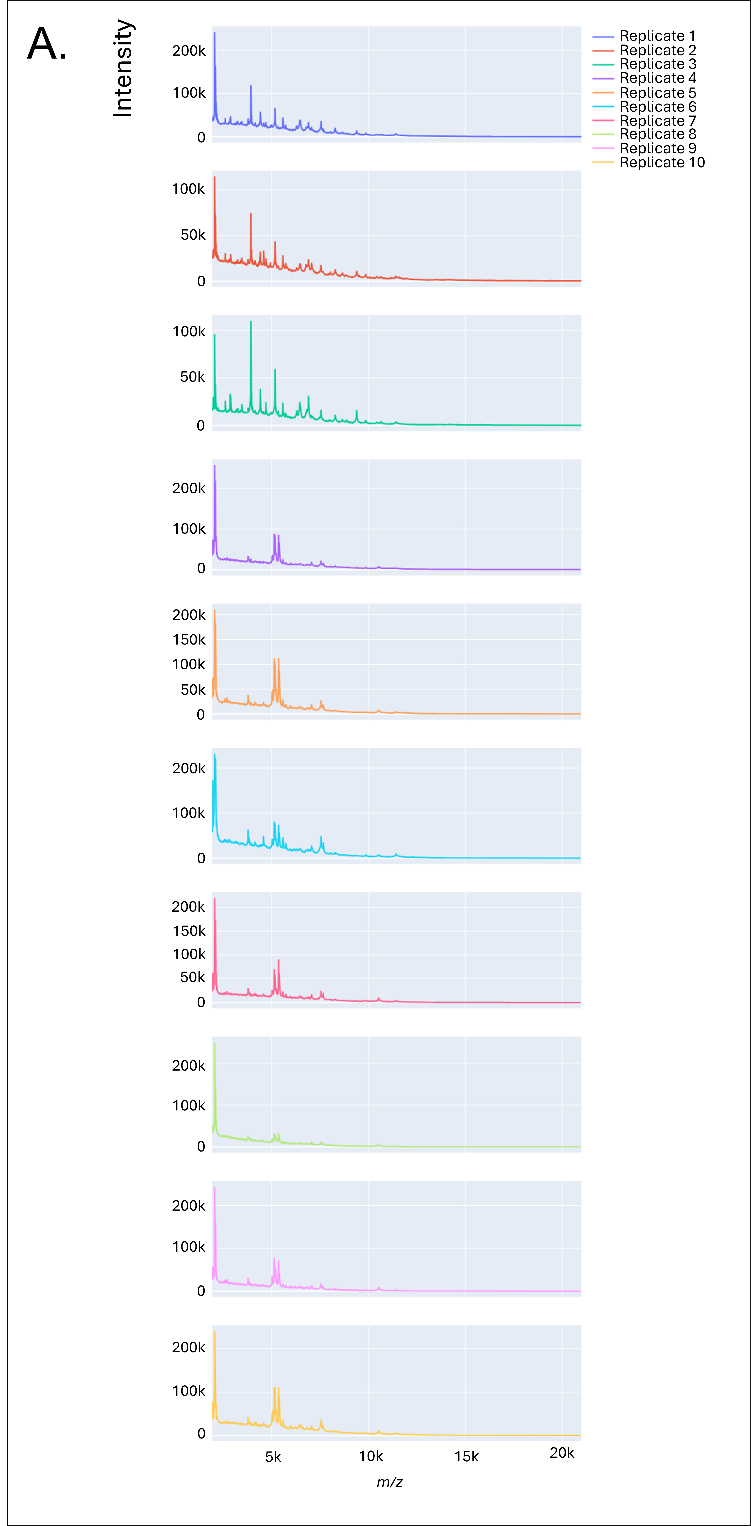

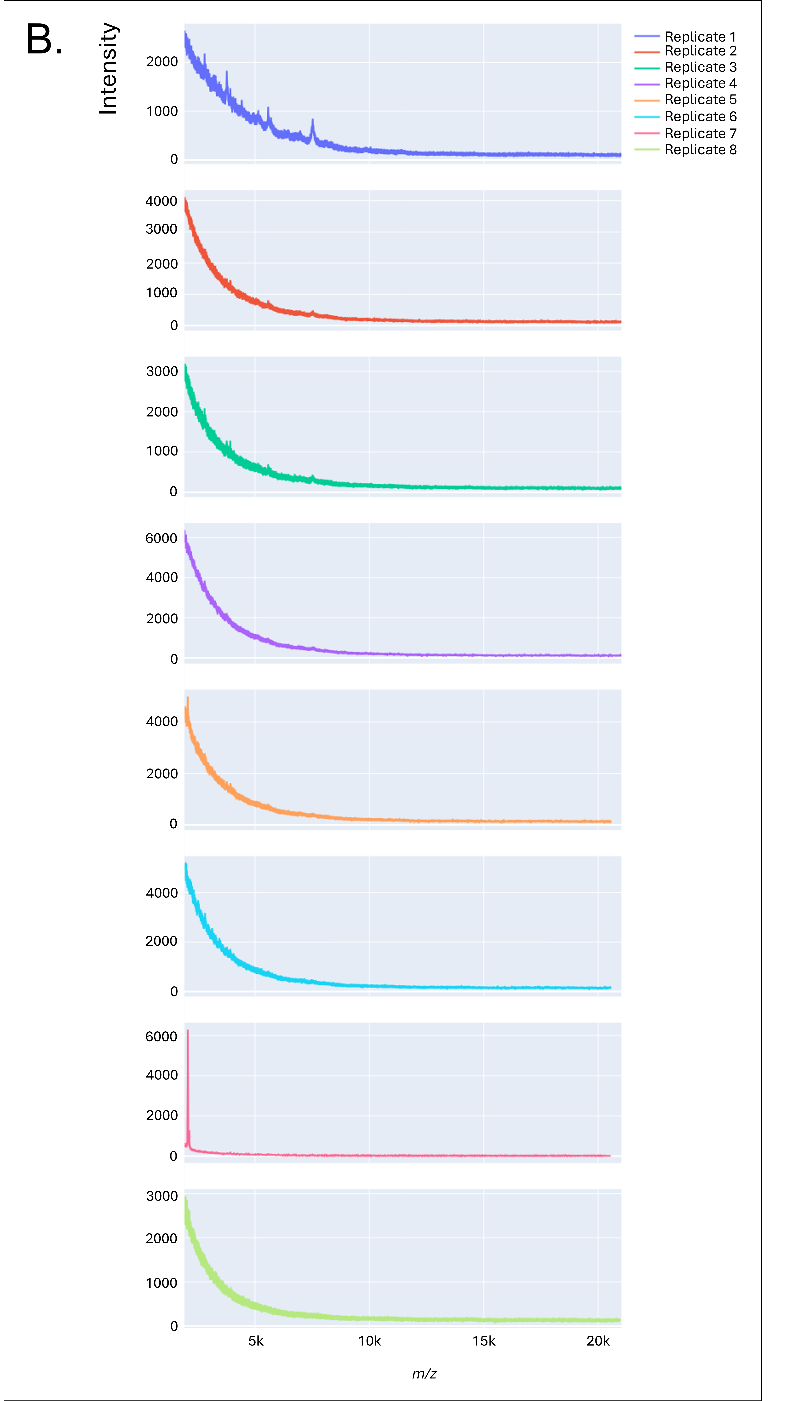

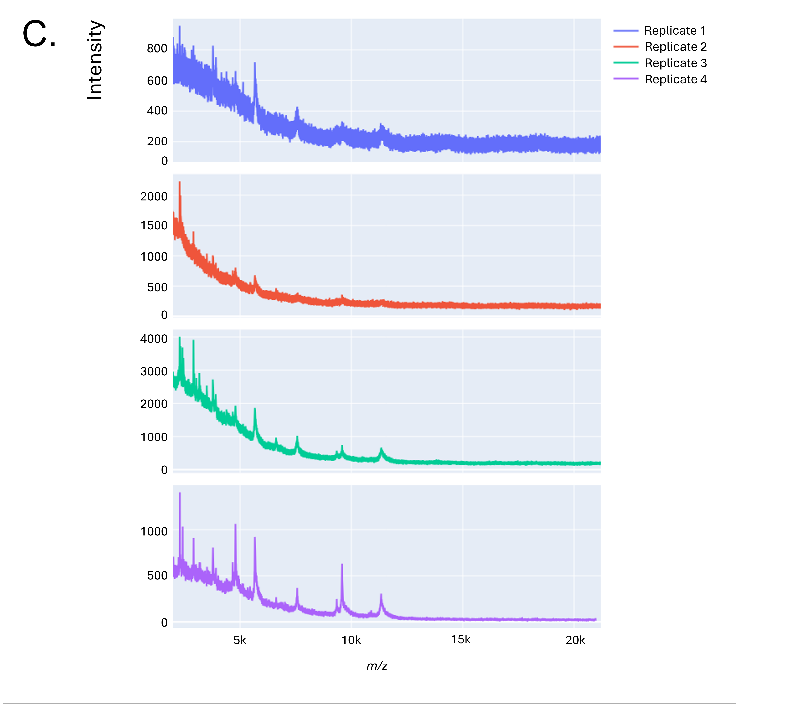
